# Fungal hydrophobins unleashed from food waste: production, rodlet assembly, and functional properties in *Ganoderma adspersum*

**DOI:** 10.64898/2026.01.25.700568

**Authors:** Carolina Reyes, Enrico Boschi, Peter Gehrig, Siiri Bienz, Naresh Kumar, Tiago Carvalho, Nico Kummer, Lukas Bürgi, Ashutosh Sinha, Francis WMR Schwarze, Silvia Campioni, Gustav Nyström

## Abstract

*Ganoderma adspersum* is a white-rot fungus (WRF) that produces amphipathic, surface-active proteins, known as hydrophobins. To explore sustainable routes for protein production and waste valorization, *G. adspersum* was cultivated under defined shaking conditions using different carbon and nitrogen sources, including apple skin food waste. From these cultures, we report for the first time the isolation and characterization of a novel class I hydrophobin, designated as Gad1. Atomic force microscopy revealed abundant rodlet-like nanostructures consistent with class I hydrophobin assemblies, and Raman spectroscopy confirmed β-sheet enrichment typical of amyloid-like organization. MALDI-TOF mass spectrometry further identified Gad1 along with additional hydrophobin-like proteins in the foam. Hydrophobin-enriched foam extracts were used to form stable oil-in-water emulsions that could be converted into porous, freeze-dried composite aerogels. These findings expand the known diversity of hydrophobins in white-rot fungi and demonstrate that food-waste-derived substrates can support hydrophobin production and functional biomaterial formation.

## Introduction

Hydrophobins are amphiphilic proteins secreted by fungi. These proteins allow fungal hyphae to breach the air-water or oil-water interface by lowering surface tension at those boundaries (1, 2). Hydrophobins also coat fruiting bodies, are used by fungi to protect their spores against wetting during dispersal(3) and help them to adhere to substrates(1). In some fungi, class I hydrophobins have been found to have a role in adhesion and pathogenic interactions(4). Once secreted, hydrophobins that encounter air or a solid surface self-assemble at the cell wall interface. This self-assembly exposes the hydrophobic amino acid residues to the environment while the hydrophilic ones interact with the polysaccharide-rich cell wall(2).

Two major classes of hydrophobins have been identified so far, class I hydrophobins and class 2 hydrophobins(5), but intermediate classes have also been described(6). Class I hydrophobins self-assemble into stable, fibrillar, and highly ordered structures that can only be disrupted by harsh acids like trifluoroacetic acid (TFA)(7) or formic acid(8). They can be isolated from foaming cultures as with SC3 from *Schizophyllum commune*(9). In contrast, class II hydrophobins form less stable structures and can be easily dissociated using milder reagents such as 60% ethanol, 2% SDS, or by applying pressure(10). Several hydrophobin class I proteins have been studied extensively and serve as models for protein structure and aggregation, including EAS from *Neurospora crassa*(11), Dew A from *Aspergillus nidulans*,(3) SC3 from *Schizophyllum commune* (12) and Vmh2 from *Pleurotus ostreatus*(13). Class I hydrophobins are small (7-20 kDa) cysteine-rich proteins with a conserved β-barrel core linked by four disulfide bridges and flexible loop regions that vary among species and affect their assembly properties(1) (14–17) .Hydrophobins are expressed at different points of the fungal life cycle(18). During its life-cycle in nature, a basidiomycete fungus will go through spore formation, monokaryotic and dikaryotic phases, primordial formation, fruiting body formation, karyogamy and finally meiosis and basidiospore formation once again(19). In *Ganoderma lucidum*, a class I hydrophobin *hyd1* was highly expressed in fungal primordia, when grown in cultivation bags, and its expression was lowest in fruiting bodies(20). Furthermore, its expression appears to be regulated by nitrogen availability and environmental stressors such as heat, salt and cell wall stress.

Food waste is an emerging resource that could potentially be used to support hydrophobin production. Globally, about one-third of all food produced is wasted, contributing significantly to greenhouse gas emissions and representing a major environmental, social and economic challenge (21, 22). Previous studies have shown that fungi and microalgae can convert food waste into secondary metabolites and proteins(23, 24). There are two studies that describe hydrophobin-like proteins isolated from food waste in fungal fermentations using *Penicillium* species and *P. ostreatus* (25, 26). However, to our knowledge, no study has yet investigated the use of *Ganoderma adspersum* to produce hydrophobins from food waste.

In this study we examined the production of class I hydrophobins by *Ganoderma adspersum* using two growth conditions. In the first, *G. adspersum* was cultured in minimal medium under shaking conditions to promote the formation of primary foam. In the second, glucose was replaced with food waste as a carbon source (**Figure 1**). Foams were collected from each condition, and following acid treatment, total protein extracts were analyzed using a range of biochemical and analytical techniques. The extracted hydrophobins were furthermore utilized to stabilize oil-in-water emulsions and to template composite cellulose-protein biohybrid aerogels illustrating their application in bio-based functional materials.

**Figure 1.**
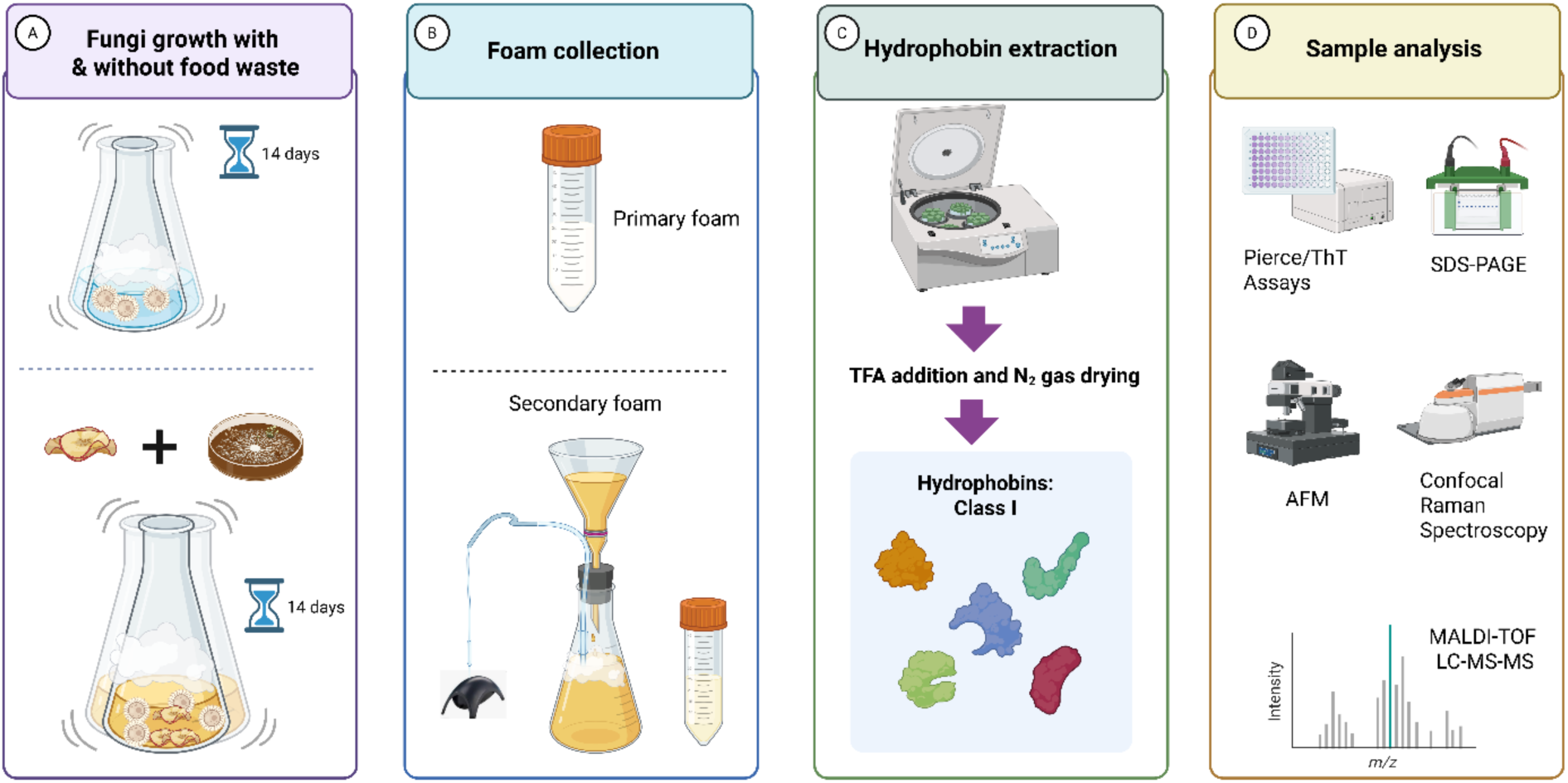
Schematic depicting the experimental approach for hydrophobin production and analysis. **A)** *G. adspersum* was grown in the absence or presence of food waste for 2 weeks in liquid media with shaking to induce primary foam production. **B)** Primary foam was collected, the remaining media was then filtered and aerated with a pump to produce a secondary foam and this foam was collected too. **C)** Class I hydrophobins were extracted from the foam by treating it with TFA. Samples were dried under N_2_ gas. **D)** Following solubilization in ethanol, samples were analyzed for total protein concentration using the Pierce and ThT Assays, SDS-PAGE, AFM, Confocal Raman Spectroscopy and MALDI-TOF LC-MS-MS.

## Results and Discussion

### Growth on different carbon and nitrogen sources

In order to assess the influence of different carbon sources, growth tests on agar plates were performed, and the diameter of mycelial growth was measured. *G. adspersum* was able to grow best in Petri dishes with minimal media when supplemented with glucose, fructose or xylose sugars. However, its growth slowed down when grown with galactose or arabinose sugars (**Figure 2A**). *G. adspersum* likely gains more energy in the form of ATP when aerobically metabolizing glucose, fructose and xylose sugars present in the minimal medium. In fungi, ATP is produced via the pentose phosphate, glycolysis and tricarboxylic acid cycle (TCA) pathways (27). However, depending on the sugar, it will enter a specific pathway, different steps will be involved in processing those sugars and therefore different amounts of ATP generated(28). Glucose and fructose would both enter the glycolysis and TCA cycles producing ∼30-32 ATP per molecule. However, xylose would first enter the pentose phosphate pathway and then glycolysis pathway generating less ATP due to the different conversion steps involved (∼5 ATP)(28). Another factor that could influence the preferential uptake of sugars in *G. adspersum* are the presence of sugar transporters. Previously, the genome of *G. adspersum* was sequenced and annotated which could provide insight about possible genes involved in various metabolic pathways (In Press). Looking into the genome of *G. adspersum*, various transporters for glucose, galactose and xylose uptake are present (**Table S1**) that could also influence which sugar is more readily taken up by *G. adspersum*. Although arabinose related genes are present, including those for export, no uptake genes were annotated in the genome (**Table S2**).

**Figure 2.**
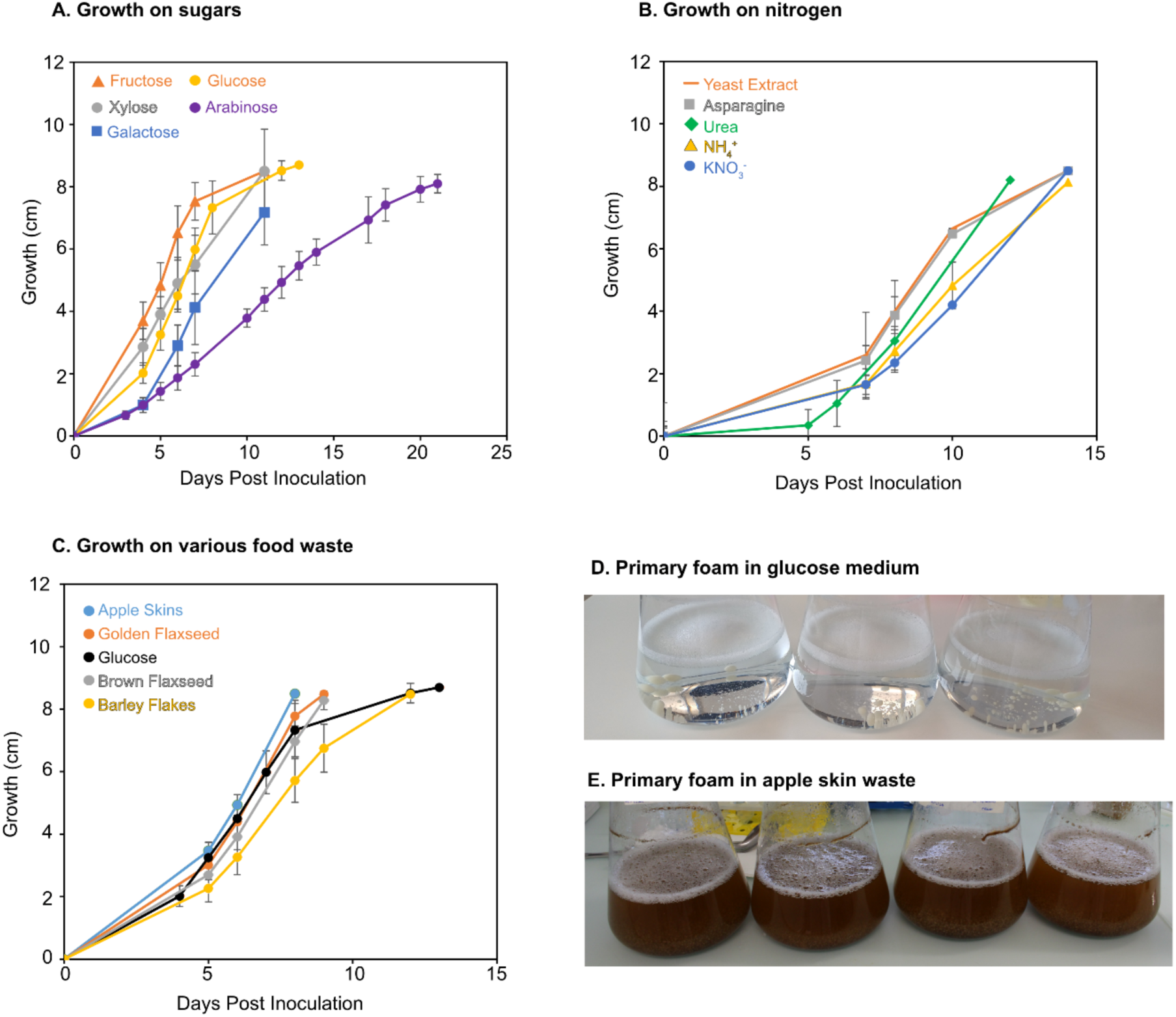
Growth of *G. adspersum* under different conditions and primary foam production. **A)** Petridish growth on different sugars in minimal media containing L-asparagine as nitrogen source. **B)** Petri dish growth on different nitrogen sources in minimal media containing glucose as carbon source. **C)** Petri dish growth on different food wastes. The media lacked glucose when grown with apple-skins and contained L-asparagine. The media contained glucose when grown with golden or brown flaxseed or barley flakes but lacked L-asparagine. **D)** Primary foam production in minimal media containing glucose. **E)** Primary foam production in minimal media containing apple skin waste.

The types of sugar that a fungus preferentially metabolizes could also depend on the types of cell structures built from these sugars. For example, the fungal cell wall and extracellular polymeric substance (EPS) is decorated with different sugar moieties like mannose and glucans (**Figure S1**). Previous studies found that the fungal culture media composition influenced the types of carbohydrates present in the EPS of *G. lucidium*(29, 30). Furthermore, when the conditions changed, including the type of sugar, so did the composition of the carbohydrates in the EPS.

Besides carbon, *G. adspersum* also grew well when yeast extract, asparagine or urea were used as an organic nitrogen source, but growth slowed down when ammonia sulphate or potassium nitrate were used (**Figure 2B**). In a gene expression study with *G. boninense*, the type of nitrogen used in the media (e.g., ammonium sulfate and potassium nitrate) was shown to either stimulate or repress the manganese peroxidase and laccase activities of *G. boninense* (31). Perhaps *G. adspersum* cannot efficiently use ammonium sulfate and potassium nitrate for enzyme synthesis and therefore grows more slowly on these substrates. Alternatively, the concentration of these two nitrogen substrates used in our media could have inhibited the growth of *G. adspersum*. In a study with *G. lucidum*, decreasing concentrations of ammonium phosphate in agitated media led to improved mycelial growth, whereas higher concentrations had the opposite effect(32).

Adding different food wastes to the growth medium instead of a supplementary sugar or nitrogen source also affected the growth of *G. adspersum*. We used carbon and nitrogen content of the food waste as a guide to determine if the food waste could replace glucose or Lasparagine in the media. Elemental analysis results showed apple skin contained the lowest amount of nitrogen compared to golden or brown flaxseeds and barley flakes (**Table S3**). Compared to the baseline SCMM, the dry food waste contained higher carbon and comparable nitrogen (**Table S3**). Therefore, we opted to use either apple skin as a substitute for glucose in the L-asparagine containing media, or the other food wastes as a substitute for L-asparagine in the glucose containing media. Results showed that *G. adspersum* grew faster on apple skin waste and flaxseed waste than on barley flakes (**Figure 2C**). It is likely that *G. adspersum* benefits from the food waste because it contains various types of carbon, including hemicellulose and cellulose(33), sugars including glucose, fructose and sucrose(34–36) and proteins as nitrogen source(37). Based on these results, we proceeded to test foam production in liquid media containing either apple skin waste (as carbon source) or flaxseed (as nitrogen source) to determine if the type of substrate would also influence foam and by extension hydrophobin production.

### Foam production on various carbon and nitrogen sources

When grown in liquid media under shaking conditions, primary foam production was observed in the minimal media (**Figure 2D**). Total protein yield was highest in the primary foam collected from minimal medium with L-asparagine (3.63 ± 1.5 mg/L) or minimal medium with yeast extract (3.58 ± 2.5 mg/L) (**Figure S2**). In contrast, when grown with inorganic nitrogen like urea or potassium nitrate, the total protein yield was lower (<0.1-1.9 mg/L) (**Figure S2**). Thus, the highest amounts of total protein in the primary foam developed in the presence of organic based nitrogen sources. Pooling of triplicate extracts from the minimal medium with L-asparagine helped to increase the total protein yield (7.5 ± 0.36 up to 16.7 mg/L) (**Figure S2**). Secondary foam production was not observed under these growth conditions. We hypothesize that this may result from a decreased synthesis or secretion of foam-stabilizing biomolecules, such as glycoproteins, when *G. adspersum* is grown in a defined minimal medium.

Primary foam production also formed in shaken cultures of *G. adspersum* containing apple skin waste **(Figure 2D**). Initially, apple skins were used to replace glucose in the minimal media with L-asparagine serving as the nitrogen source. Despite L-asparagine being present, *G. adspersum* produced less protein (0.18 mg/L) than in the minimal media with no food waste (3.63 ± 1.5 mg/L) (**Figures S2 and S3**). Apple skins contain lignocellulose materials and sugars (38, 39). However, these polysaccharides must first be enzymatically degraded before the fungus can access the sugars. Although *G. adspersum* showed normal growth in the cultures (**Figure 2C**), the lower protein yield in the foam may indicate a shift in metabolism rather than carbon limitation as such. It is possible that, under these conditions, *G. adspersum* allocates resources toward the production of different extracellular proteins or enzymes involved in substrate degradation instead of foam-associated proteins such as hydrophobins. Replacing L-asparagine with yeast extract in the media, increased the pooled protein concentration (up to 7.4 mg/L) in the primary foam samples (**Figure S3**). Yeast extract is rich in free amino acids (30-40%), including glutamic acid, and contains some sugars (40). It could be that the yeast extract provides *G. adspersum* with a more diverse cocktail of amino acids compared to L-asparagine alone leading to more protein production in the foam. Previous studies found yeast extract to be an excellent organic nitrogen source increasing enzyme production in white-fungi cultures (41, 42). In addition, apple skins alone (no glucose or L-asparagine) did not produce any foam in control experiments (**Figure S4**), confirming that apple skins by themselves are insufficient to induce foam formation. Notably, fungal growth was still observed under these conditions (**Figure 2C**), suggesting that growth rate is not necessarily coupled to the production of extracellular or foam-associated proteins. It is also important to note that foaming phenomena are specific to aerated or shaken liquid cultures, where hydrophobins accumulate at the air-liquid interface, and may not be directly comparable to static growth observed on agar media.

Unlike the minimal media experiments above, secondary foam formed in the apple skin containing samples when aerated with an aquarium pump. The highest protein yield was obtained when secondary foam replicate samples from apple skin waste and yeast extract were pooled together (16 mg/L) (**Figure S3**). Perhaps the growth conditions with the apple skin waste and yeast extract result in higher production of foam stabilizing compounds such as glycoproteins that are released by the fungus. Previous studies with yeast have found that they release glycoproteins, such as mannoproteins from their cell walls, that are excellent foam stabilizers, during alcohol fermentation of beer and wine(43, 44).

We also tested the possibility of using flaxseed (golden and brown) and barley flakes, as an additional substrate. However, almost no foam was produced after 11 and 12 days of incubation (**Figure S5**). Despite the flaxseeds having a growth promotion effect on solid media containing apple skin waste (**Figure S6**), it did not promote hydrophobin production under shake flask conditions. It could be that the higher nitrogen content of the flaxseed and barley flakes (**Table S1**) negatively regulate hydrophobin production, preventing its formation under shake flask conditions. Previous studies in *G. lucidium* (a related species) found that the expression of the hydrophobin gene *hyd1* increased under nitrogen limiting conditions due to negative regulation of this gene by the transcriptional regulator AreA(20). This transcriptional regulator is typically involved in regulating genes involved in nitrogen metabolism in filamentous fungi by either promoting the expression of nitrogen genes or repressing them(45, 46). More detailed expression-based experiments, like the study above, would further clarify if and how nitrogen has a role in regulating the expression of hydrophobin genes in *G. adspersum*.

### MALDI-TOF mass spectrometry of hydrophobin proteins

Samples analyzed by SDS-PAGE showed the presence of an ∼ 10 kDa protein signal in total protein extracts from both the minimal medium and apple skin waste medium (**Figure 3A and B**). Because of their small size, we interpreted the bands as hydrophobins. Comparison of samples before and after reduction showed that when reduced, the mass of the sample shifted by 8 Da (**Figure 3C and D**). In previous studies, hydrophobins were reduced with dithiothreitol (DTT) prior to MALDI-TOF analysis to break disulfide bonds and improve ionization of the small cysteine-rich proteins (47). In our analysis, samples were instead reduced using tris(2carboxyethyl)phosphine (TCEP) in tetraethylammonium bromide. TCEP performs the same reductive function as DTT but is more stable, odor-free, and effective under neutral to mildly basic conditions, making it compatible with downstream MS sample preparation. Similarly, mass spectrometry analysis of samples from food waste cultivation also revealed the presence of hydrophobins in secondary foam samples based on the initial mass of the proteins prior to reduction (**Figure S7**).

**Figure 3.**
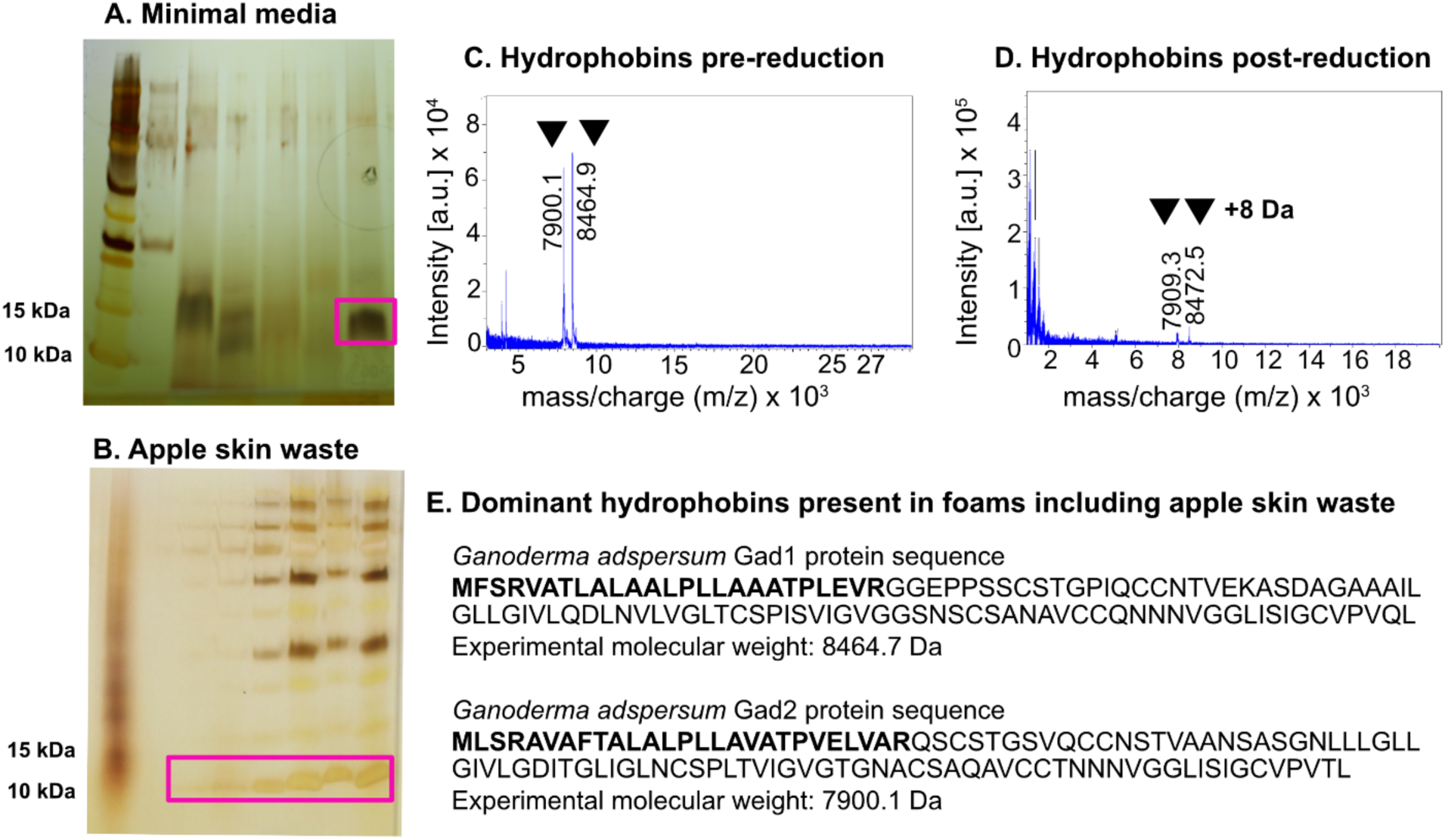
Identification of class I hydrophobins in the primary foam of *G. adpsersum* when grown on minimal media or media with apple skin waste. **A and B)** SDS-PAGE gels of small proteins in the size range of hydrophobins highlighted with a box. **C)** Mass spectra showing the mass distribution of hydrophobins present in the minimal media before the disulfide bonds between cysteine residues were reduced **D)** The shift in mass of the same samples in C showing how reduction of the disulfide bonds leads to an increase of mass by 8Da when hydrophobins are present. **E**) Hydrophobins that were unambiguously present in the samples. The predicted export signal sequence is in bold.

### Identification of hydrophobin proteins

MALDI-TOF analysis revealed that one hydrophobin in particular comprised the sample, here called Gad1 (**Figure 3D**).

Removal of the N-terminal sequence, during *in silico* analysis, was required to correctly identify the hydrophobin, as observed in pervious mass spectrometry studies of HFBI(48) and HFB2(49), which also show the presence of an N-terminal sequence. It is hypothesized that this N-terminal sequence is post-translationally cleaved prior to secretion of hydrophobins out of the cell (48, 49). Thus, the measured mass of the hydrophobin is lower than would be calculated based on its amino acid sequence alone. Another hydrophobin ∼7.9 kDa was also observed in both culture types designated Gad2 (**Figure 3E**). During *in silico* analysis, removal of its Nterminal sequence, was also needed to correctly identify this protein. Furthermore, alignment of Gad1 and Gad2 with other class I hydrophobin amino acid sequences revealed a high similarity with respect to the eight cysteine residues (**Figure S8**).

### AFM images of rod-like structures present in minimal growth medium

To further confirm that the two hydrophobins identified in *G. adspersum* had the self-assembled morphology typical of class I hydrophobin, AFM imaging was performed. AFM results showed the presence of short rod-like structures in total protein extracts from primary foam of *G. adspersum* grown on minimal medium (**Figure 4**). These short rods appeared stacked on top of each other to form larger rod-like structures. Similar rod-like structures have been observed in *Schizophyllum commune*(50), *Neurospora crassa*(11), *Aspergillus nidulans*(51), and *Pleurotus ostreatus* (52) by AFM analysis. Class I hydrophobins are known to form stable monolayers and latterly arranged fibrils (51). Thus, these results were a first confirmation that class I hydrophobins were present in the primary foam. These rod-like structures also appeared in primary and secondary foam samples from apple skin cultures. They were also stacked on top of each other similar to the rods observed in the minimal media (**Figure 5A-C**). Closer imaging also revealed a branching order to the stacked rods similar to what has been observed for cellulose nanofibrils (53) or amyloid fibrils (54)) (**Figure 5C-E**). Sometimes the presence of round particles was observed but it was unclear if this was protein or particles from the food waste extraction. Rodlet contour lengths were quantified from an AFM image (**Figure 5A**) of apple skin derived sample using FiberApp (n=190 fibers)(55). The rodlets displayed an average contour length of 211 ± 68 nm, with most fibers ranging between ∼140 and 280 nm and a small proportion extending up to ∼450 nm (**Figure S9**). These measurements confirm that the rodlets assemble into heterogeneous but consistently nanoscale fibers, in agreement with previously reported ranges for class I hydrophobins(50).

**Figure 4.**
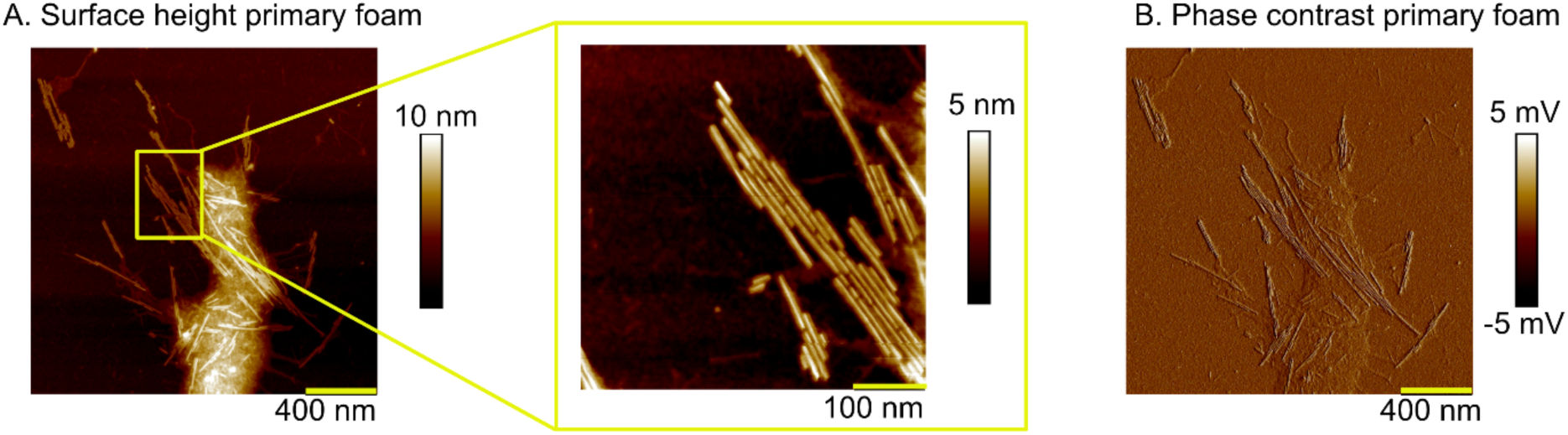
AFM visualization of rodlets from the primary foam of *G. adspersum* when grown on minimal medium. **A**) Surface height of primary foam sample and close-up section. **B**) Phase contrast of primary foam sample.

**Figure 5.**
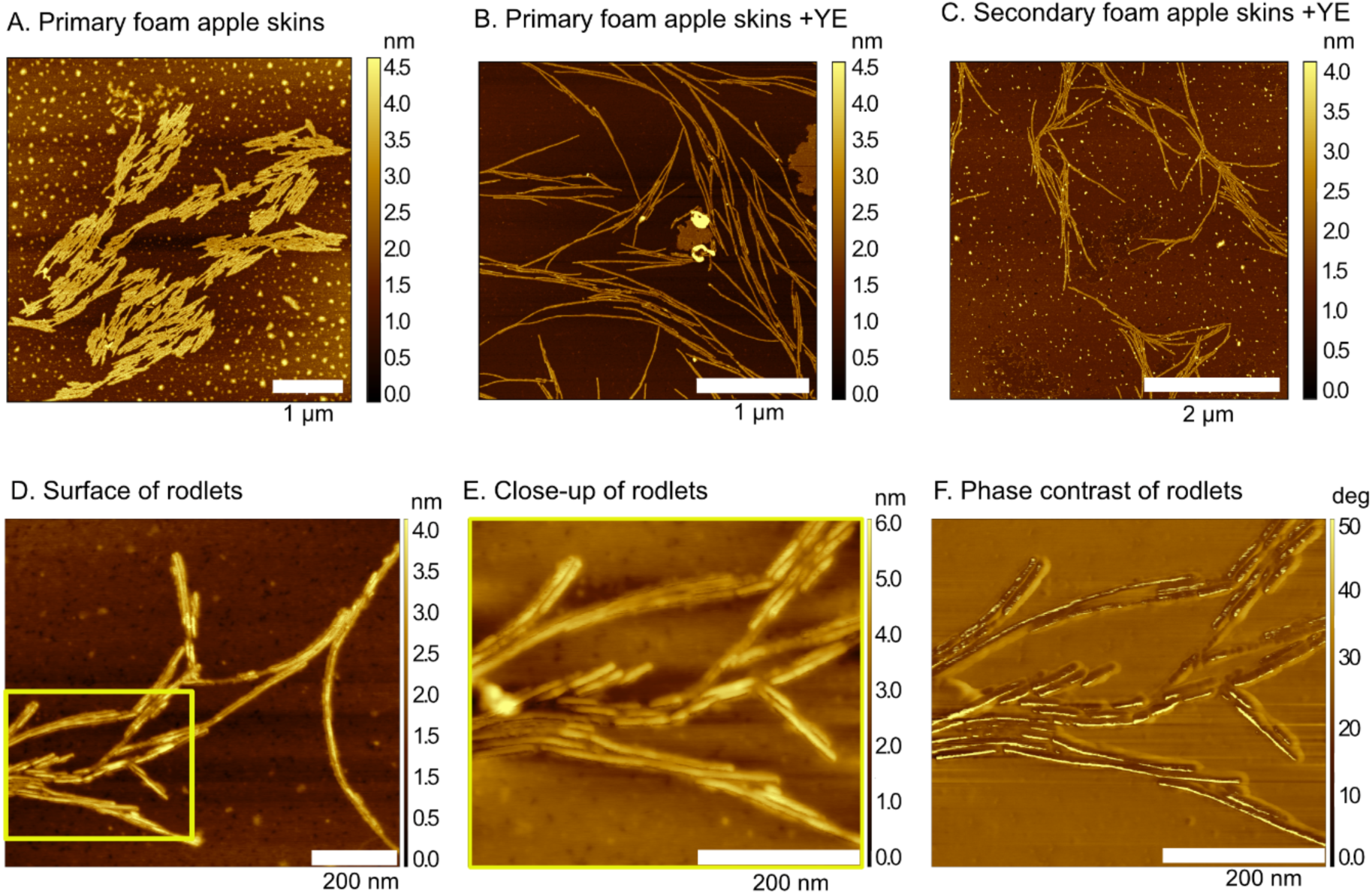
AFM visualization of rodlets from the primary foam of *G. adspersum* when grown on apple skin waste. **A)** Rodlets produced in primary foam when *G. adspersum* is grown with apple skins in minimal media lacking glucose. **B)** Rodlets produced in primary foam and in **C)** secondary foam, when *G. adspersum* is grown with apple skins and yeast extract. **D)** Close up of the surface of the rodlets from secondary foam **E)** Close-up of the rodlets. **F)** Phase contrast image of the rodlets in E.

### Confocal Raman spectroscopy

In confocal Raman spectroscopy, the Amide I band (1600-1700 cm-1) is widely used to assess protein secondary structure, as it primarily reflects C=O stretching of the peptide backbone (∼80%), with minor C-N and N-H contributions(56). The Amide I band is highly sensitive to hydrogen bonding and molecular conformation enabling the distinction of α-helices, β-sheets, turns, and unordered structures by Gaussian deconvolution of the overlapping Raman signals. Raman spectra of the three protein aggregates from the rodlet sample derived from primary foam (non-food waste) displayed consistent vibrational signatures, demonstrating that on average protein rodlets adopt a largely uniform structural organization (**Figure 6A**). Amide I deconvolution yielded three peaks 1666 cm^−1^ (sharp dominant, 66.0 ± 0.5%), 1650.5 cm^−1^ (broad; 29.4 ± 0.3%), and 1678.5 cm^−1^ (sharp; 4.4 ± 0.8%) (**Figure S10**). The 1666 cm^−1^ peak is indicative of parallel β-sheets or turns, consistent with highly ordered amyloid rodlets(57). The 1650.5 cm^−1^ band reflects alpha-helix structures(58). The 1678.5 cm^−1^ peak corresponds to coil or polyproline II (58). Together, these data indicate a β-sheet-rich architecture with alpha helices and minor coil components. Raman spectra result of three different rodlet samples derived from secondary foam (apple skin waste), also showed similar vibrational characteristics (**Figure 6B**). Just as with the sample derived from the defined medium (no apple skins), Amide I deconvolution yielded three peaks at 1666 cm-1, 1650.5 cm-1 and 1678.5 cm-1 (**Figure 6C**). Together, these data also indicate a β-sheet-rich architecture (65.0 ± 0.4%) with alpha helices (28.8 ± 0.3%) and minor coil (5.6 ± 0.4%) components (**Figure 6D**). When compared to FTIR secondary structure information from the model class I hydrophobin SC3, formed at an air/water interface, the rodlet secondary structural composition is similar (**Table S4**). Perhaps *G. adspersum* rodlet formation occurs similarly to SC3. During SC3 rodlet formation, there are three different rodlet states, a monomeric amorphous state, an intermediate state that is mostly rich in alpha helices and two beta sheet states, one of which is more stable than the other(59). To obtain a more definitive answer, class I hydrophobins Gad1 and Gad2 would have to be purified and further characterized as in the above study with SC3.

**Figure 6.**
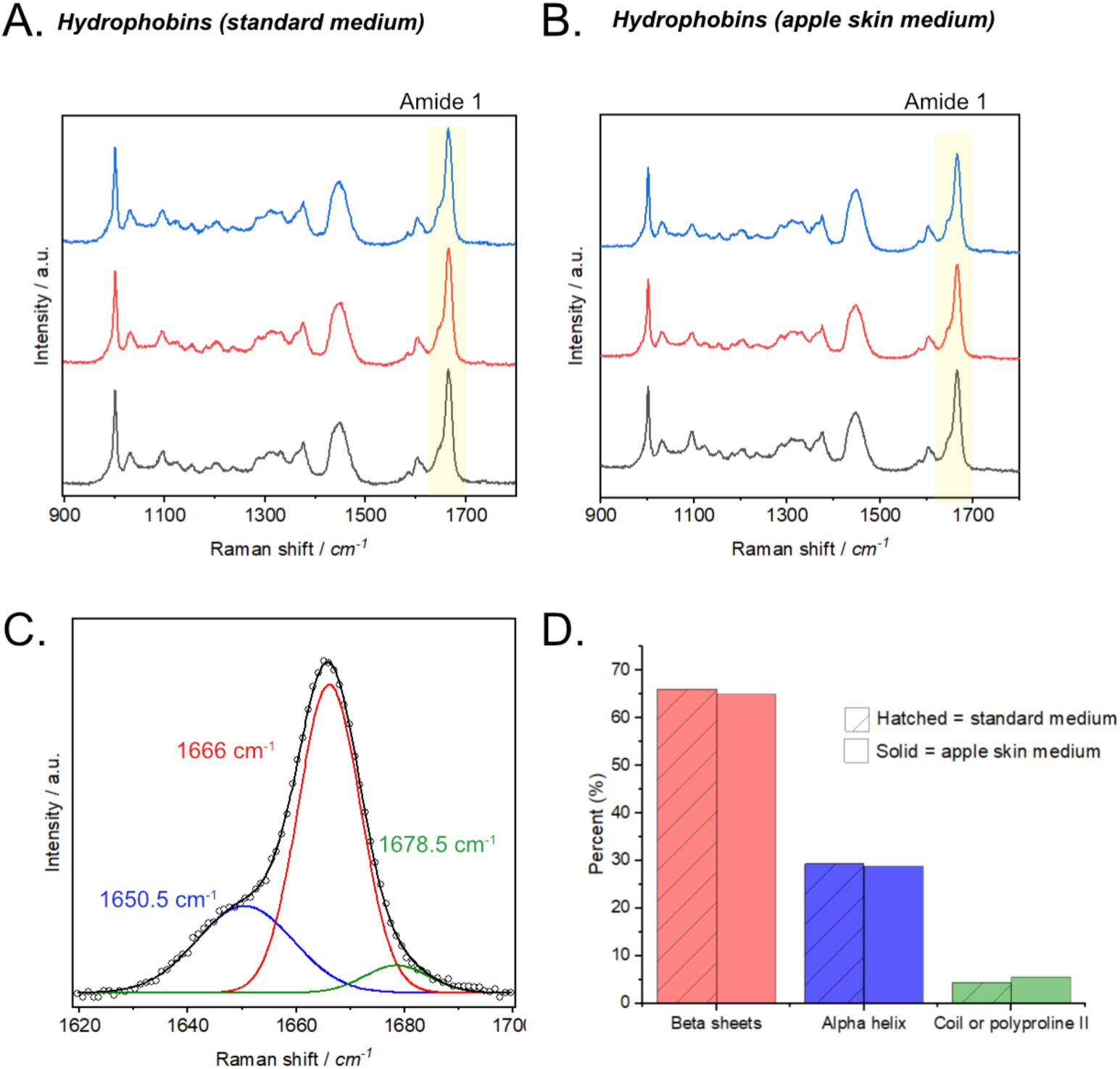
Confocal Raman Spectra of Class I Hydrophobin Protein Aggregates. **A)** Raman spectra of hydrophobin aggregates, from foams produced in non-food waste medium. The Amide I region is highlighted. **B)** Raman spectra of one of the hydrophobin aggregates, from foams produced in apple skin (food-waste) medium. The Amide I region is highlighted. **C)** Representative deconvolution of the Amide I band, from foams produced in apple skin (food-waste) medium, showing Gaussian fitting into β-sheet, α-helix, and coil/polyproline II components **D)** Relative secondary structure contributions from Gaussian fitting of spectra in A and B, showing β-sheets as the dominant feature under both cultivation conditions.

### Thioflavin T assay

Thioflavin T (ThT) binds to amyloid-like fibrils formed by class I hydrophobins through their beta-core structure, allowing relative quantification of these proteins. Previous studies have shown that as the concentration of hydrophobins or amylogenic proteins increases, rodlet/fibril formation also increases, leading to higher ThT fluorescence (60, 61). We applied the ThT assay to estimate the amount of class I hydrophobins in total protein extracts obtained from foam samples. A ≥ 3-fold increase in fluorescence relative to the non-protein control was considered a positive indication of hydrophobin presence, consistent with previous reports that a 2-5 increase in ThT signal reflects amyloid-like fibril formation (62, 63).

Secondary foam samples derived from apple skin waste showed higher fluorescence compared to primary foam samples from non-food waste (**Figure S11**). This increase occurred despite similar total protein concentrations and, notably, stronger hydrophobin bands in the SDS-PAGE of minimal-media foams. The higher ThT signal in the apple-skin samples therefore likely reflects differences in the composition or structural state of the secreted proteins rather than a greater abundance of hydrophobins. It is possible that additional proteins present in the apple-skin foams contribute to ThT binding, or that *G. adspersum* produces alternative hydrophobin variants or aggregated forms of Gad1/Gad2 that persist after TFA treatment. Thus, while ThT fluorescence indicates enhanced amyloid-like assembly under these conditions, it does not necessarily correspond to higher hydrophobin content.

Because ThT can bind to beta-sheet fibrils in general, we verified the molecular origin of the signal. Raman spectroscopy of the secondary foam extracts confirmed beta sheet enrichment, and AFM imaging revealed rodlet-like nanostructures consistent with class I hydrophobin assemblies. Foam protein extracts were disassembled with TFA and then reconstituted in phosphate buffer, after 30 min, ThT signal was robust, indicating that the signal depends on the self-assembled state of the protein under our assay conditions. Moreover, LC/MS analysis identified two class I hydrophobins as the major proteins in secondary foam. Together, these data support that hydrophobin rodlets are the primary source of the ThT fluorescence in our system.

Comparable findings in *Pleurotus ostreatus* support a functional role for hydrophobins in substrate degradation and the hydrophobin gene *vmh3* was specifically induced when lignin served as a growth substrate (64, 65). In our shaken liquid cultures, the role of hydrophobins is unlikely to be substrate “attachment” in the classical sense. Instead, apple skins are rich in hydrophobin cuticular waxes (long-chain alkanes, fatty acids, esters, triterpenoids) and cutin, which remain hydrophobic even after autoclaving, though their physical organization may become less crystalline(66). Hydrophobins likely accumulate at these hydrophobic interfaces, increasing wettability and dispersal of wax-rich fragments in the medium. This could in turn improve the accessibility of apple skin components to degradative enzymes such as cellulases, hemicellulases, and lignin-modifying oxidoreductases (e.g. laccases, manganese peroxidases, versatile peroxidases) (67). By analogy to the *P. ostreatus* studies, the elevated hydrophobin levels observed in *G. adspersum* foams may therefore reflect a functional role in overcoming the recalcitrant cuticle of apple skins and facilitating their degradation.

### Porous Freeze-Dried Emulsion Composites (FDECs)

As a proof of concept, hydrophobin-enriched secondary foam derived from apple skin waste was used to make oil-in-water emulsions and porous freeze-dried emulsion composites (FDECs). Emulsions containing coconut oil, phosphate buffer, and TEMPO-oxidized cellulose nanofibers were stabilized by hydrophobins and remained intact for ≥ 30 days. SDS-stabilized emulsions also showed stability over the same period, whereas controls lacking hydrophobins partitioned after 12 days (**Figure 7A**). Using hydrophobin samples we also fabricated FDECs that retained stable foam architectures comparable to controls (**Figure 7B**). SEM imaging revealed that hydrophobin-containing FDECs formed highly porous open networks with large, interconnected cavities (**Figure 7C**). By contrast, non-hydrophobin control showed smaller, more compact pores with thicker walls and a denser morphology (**Figure 7C**).

**Figure 7.**
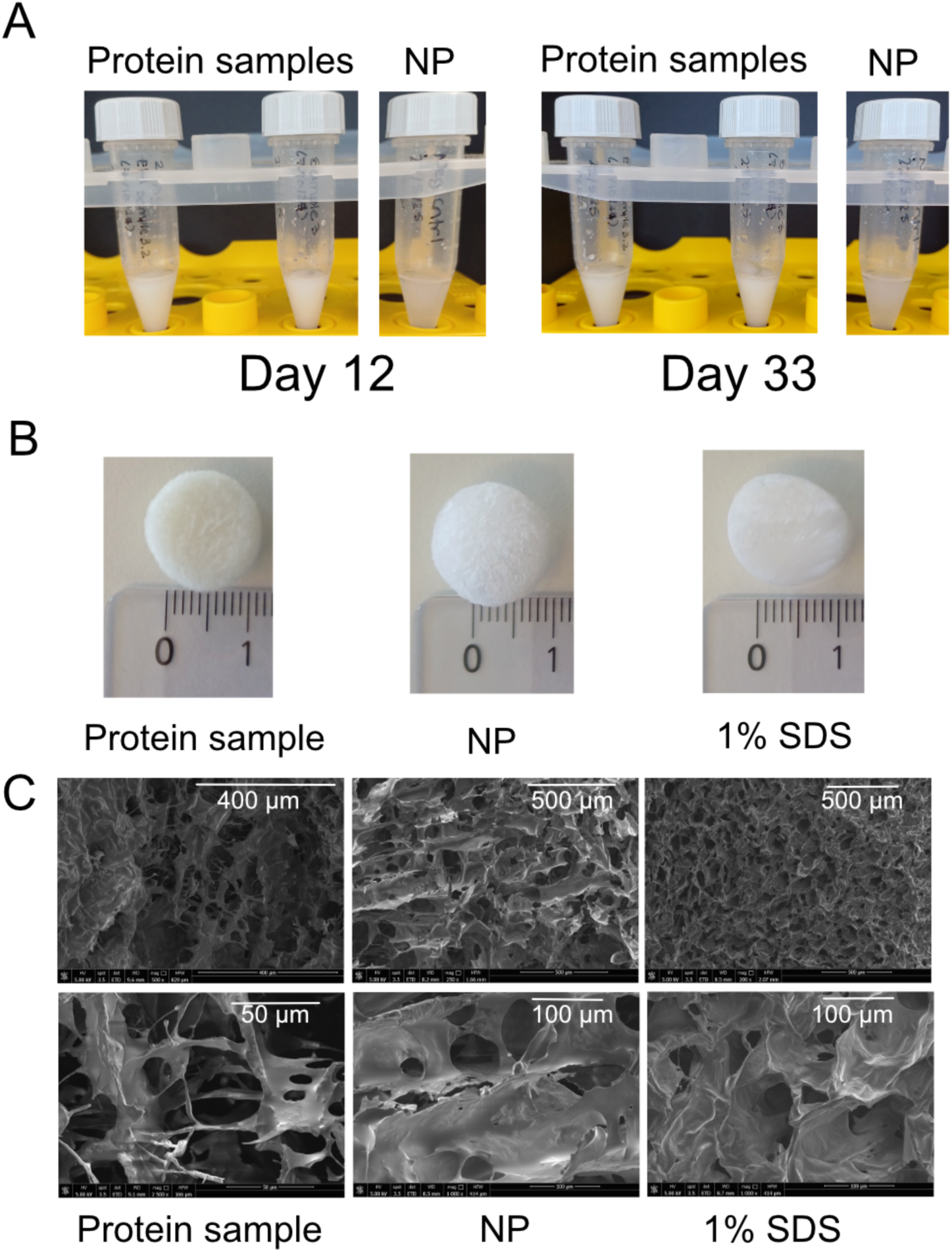
Emulsions and porous freeze-dried emulsion composites (FDECs) prepared with hydrophobins. (A) Oil-in-water emulsions containing coconut oil, phosphate buffer, and TEMPO-oxidized cellulose nanofibers stabilized with hydrophobins remained intact for ≥30 days. Controls lacking hydrophobins were less creamy and more transparent in comparison at day 12. NP=no protein control (B) Hydrophobin-containing emulsions were freeze-dried to produce FDECs with stable foam architectures, similar to controls. (C) SEM images of TEMPO-CNF FDEC cross-sections containing hydrophobins (Protein Sample), CNF-only control (NP), and SDS control (1% SDS). The hydrophobin-containing composites displays a more heterogeneous pore network with thinner, textured lamellae and occasional fibrillar bridges between pores, features consistent with partial adsorption or aggregation of hydrophobin-like proteins on the CNF matrix. In contrast, the CNF-only control (NP) exhibits smoother and more open pores typical of unmodified CNF aerogels, while the SDS control shows a denser and more compact structure with reduced porosity. Scale bars are indicated in each image.

## Conclusions

This study demonstrates, for the first time, that food waste can serve as a viable substrate to produce class I hydrophobins using *G. adspersum*. Apple skin waste supported the formation of both primary and secondary foams containing hydrophobin-like proteins that retained their characteristic amyloid-like properties. The ability to obtain functional, surface-active proteins from a lignocellulosic waste stream highlights a sustainable route for valorizing food by-products. Although the total extractable protein quantities remain low, these hydrophobin-rich foams exhibited notable interfacial and emulsifying capabilities, indicating that even small amounts can exert strong surface activity. Future studies could focus on enhancing hydrophobin production, such as through overexpression of *gad1* and *gad2*, and on exploiting these bioderived foams as functional components in sustainable emulsions, coatings, and film-forming materials.

## Materials and Methods

### Strain information and growth conditions

*Ganoderma adspersum* CBS 147723 was grown on 2% malt extract agar (MEA) Petri dishes (prepared using distilled water) and incubated at 30°C and 40% relative humidity (RH) and their growth tracked by measuring cm of growth until the whole Petri-dish was occupied with fungal growth. To determine if *G. adspersum* could produce class I hydrophobins, 5 agar plugs from an MEA Petri dish were always transferred to a 2L flask containing 1 L of minimal media (prepared using distilled water): 22 g/L glucose monohydrate, 1.5 g/L L-asparagine monohydrate, 0.5g/L MgSO_4_ 7H_2_O, 1 mL trace element solution [0.06 g/L HBO_3_, 0.04 g/L (NH_4_)_6_Mo_7_O_24_ 4H_2_O, 0.2 g/L CuSO_4_ 5H_2_O, 2 g/L ZnSO_4_ 7H_2_O, 0.1 g/L MnSO_4_ 4H_2_O, 0.4g/L CoCl_2_ 6H_2_O, 1.2 g/L Ca(NO_3_)_2_ 4H_2_O], 1 mL thiamine stock solution (0.4 mM), 1 mL FeCl_3_ 6H_2_O stock solution (18 mM) and 3 mL phosphate buffer solution [1.35M KH_2_PO_4_, 3M K_2_HPO_4_]. This recipe is based on media for the hydrophobin producing white-rot fungus *Schizophyllum commune*(68). Liquid cultures were incubated at 30°C, 40% RH, and 150 rpm until foaming was observed at the top of the culture medium.

The growth of *G. adspersum* was tested on minimal media agar plates (same recipe as above plus 14g/L agar) where the glucose or L (+)-asparagine were replaced with other carbon and nitrogen sources. 0.1 M L-arabinose, D-galactose, D-fructose or D-xylose substituted glucose in some experiments. To test different sources of nitrogen (N), L-asparagine was replaced in the minimal media with 2.7 g of yeast extract or 0.01 M of urea, ammonium sulfate or potassium nitrate. Petri-dishes were incubated as described above.

For the food waste experiments, 5g of dry apple skin waste (Ramseier Suisse AG and Landi Sursee) was used as the main carbon source and liquid cultures were incubated up to 11 days before harvesting the primary foam. Golden and brown flaxseed (ETHZ) or barley flakes (ProSeed Ingredients SA) were used to substitute L-asparagine.

### Chemical characterization of the food waste

The carbon and nitrogen content of dry food waste samples were determined with a Leco TruSpec CHN micro elemental analyzer by the ETHZ Molecular and Biomolecular Analysis Service. For the analysis, dry samples were milled to a fine powder using a commercial food processor. The powder was used to fill capillary tubes for elemental analysis.

### Extraction of hydrophobins from culture media

Following cultivation (30°C for 2-2.5 weeks), class I hydrophobin accumulates/upconcentrates at the interface of the air and media as a foam defined here as primary foam. To further induce hydrophobin production, the media is first filtered using paper. Next, the filtered solution is aerated with an aquarium air pump (Tetra Tec APS 100) and an air stone for up to 20 minutes to induce further foam production, defined as secondary foam. The foam was collected using a steel spatula and placed in a 50 mL polypropylene falcon tube (VWR Ultra-high performance centrifugation tubes) and centrifuged at 10,000 rpm for 30 min at room temperature. Afterwards, the supernatant was discarded and the foam pellet washed up to 3 times with MilliQ water, sonicated with a water sonicator for 15 min in between washes if the sample contained aggregates, and centrifuged as before in between washes. After the final wash, the pellet was dissolved in 1-2 mL of concentrated trifluoroacetic acid (TFA), sonicated for 15 min in a water sonicator if the pellet was aggregating, and dried under a stream of air or N_2_.

### Protein concentration and size estimations

Total protein concentration was determined by first resuspending the dry sample in 2-5 mL of 10% ethanol, using the BCA Pierce Assay (Thermo Fischer) using the enhanced test tube method following the manufacturer’s instructions. Ten percent ethanol was used since 10% ethanol has been found to slow down aggregation of DewA class I hydrophobins (69) and could facilitate SDS-PAGE analysis and protein quantitation. Low protein binding tubes (Thermo Fisher) were used for sample preparation. To estimate protein size, an aliquot of the ethanol dissolved sample was TFA treated, dried under a stream of air or N_2_ and dissolved in MilliQ water. The sample were then combined with NuPAGE LDS Sample Buffer 4X (Thermo Fisher) and heated at 70°C for 10 min. Samples were loaded onto 4-12% NuPage Novex Bis-Tris precast gels using NuPAGE MES SDS Running Buffer (Thermo Fisher) to resolve low molecular weight proteins (2-200 kDa) under denaturing conditions. NuPage Reducing Agent was added to all samples prior to loading and NuPage Antioxidant solution (Thermo Fisher) was added to the running buffer as recommended in the user manuals. The Page Ruler Pre-stained protein ladder (Thermo Fisher) was diluted 10X using NuPAGE LDS Sample Buffer 4X and MilliQ water, and 5 μL was loaded per lane. To visualize the protein bands, the Silver Staining Express Kit (Thermo Fischer) was used following the manufacturer’s instructions.

### MALDI-TOF LC-MS/MS

A TFA treated sample was reconstituted in 40 μL of 60% ethanol prior to MALDI-TOF MS analysis. Samples were spotted onto the MALDI target (MTP 384 target polished steel TF, Bruker Daltonics, Bremen, Germany) with saturated sinapinic acid solution in acetonitrile/water (50:50) for subsequent MALDI-TOF measurements. The sample were also desalted using C18 Ziptips (Millipore). MALDI measurements were performed on an RAPIFLEX MALDITOF/TOF mass spectrometer equipped with a smartbeam laser (Bruker Daltonics). The measurement parameters were programmed in felxControl (Version 4.2.8.1): laser frequency of 2,000 Hz in the positive linear mode with acquisition ranging from 3,000 to 30,000 Da. Final spectra consisted of 4000 shots per analysis. MS data of ions of interest were acquired manually.

To determine if the sample contained potential hydrophobin proteins a reduction step was carried out using the same reconstituted sample as above based on previously published methods(47). Samples were reduced by adding tris(2-carboxyethyl)phosphine (TCEP) (5 mM) in tetraethylammonium bromide (TEAB) (50 mM) over 1 h at room temperature. Next, samples were spotted and processed for analysis as described above.

To determine the identification of the putative hydrophobins in the sample, 5 μL (∼ 25 μg) of the reconstituted sample was reduced by adding 5 μL of TCEP (10 mM) in TEAB (100 mM) over 1 h at 37°C. The sample was acidified by adding 50 μL of TFA 0.01%, resulting in a pH value < 3, and digested with pepsin A (0.5 μg in 0.1% TFA) at 37 °C overnight. The samples were dried, desalted using Stage Tips and dried again.

The samples were redissolved in 20 μL of acetonitrile (3%)/formic acid (0.1%) for the LC-MS analysis. Peptides were separated on an M-class UPLC (Waters) and analyzed on an Orbitrap Fusion Lumos Tribrid mass spectrometer (Thermo Fisher Scientific). Higher-energy collisional dissociation (HCD) spectra were acquired in the linear ion trap of the instrument.

### Proteomics data analysis

The acquired MS data were converted to the Mascot Generic file format (.mgf files) and were processed for identification using the Mascot search engine (Matrix Science). Joint Genome Institute (JGI) assembly and annotation of *Ganoderma adspersum* CBS 147723 v1.0 (https://mycocosm.jgi.doe.gov/Ganads1/Ganads1.info.html) was used for MS identification(70). The spectra were searched against the JGI filtered *G. adspersum* protein sequences (Filtered Models (“best”)) file (1_GeneCatalog_proteins_20221111.aa.fasta.gz). The following modification was set: Variable modifications [acetyl (protein N-term), oxidation (M). The four primary cleavage sites of pepsin A (F, L, Y and W) were taken into account, and semi-specific search parameter were applied. The protein identification results were imported in the Scaffold (Proteomics Software) to visualize the protein and peptide identification results. For data analysis, a false discovery rate (FDR) of 1% was used with a minimum number of peptides per protein set as 2, and a peptide FDR of 0.1%. Additionally, decoy entries (false positives) were included as a control to ensure the FDR cut-offs selected were working.

### Atomic Force Microscopy (AFM) imaging

AFM imaging was conducted using a Brucker Dimension Icon 3 equipped and Bruker RTESPA-150 probes. For AFM imaging 500 μL of hydrophobin samples (in 10% ethanol) were dried under a stream of air and the hydrophobins solubilized by TFA evaporation as described above. Low Protein Binding tubes (Thermo Fisher) were used to prepare samples. The samples were next dissolved in 500 μL 60% ethanol and diluted 10X with MilliQ water. Again, the use of ethanol has been shown to alter the aggregation of class I DewA hydrophobins by slowing this process down, while long incubation times could facilitate the formation of rodlets(69). Samples were incubated at room temperature from 2-4 hours before a 50 μL aliquot was taken and applied to the AFM slide. The slide was dried overnight inside a glass Petri-dish container prior to imaging. The images in Figure 4 were captured in tapping mode at a scan rate of 0.5 Hz and a resolution of 1024 × 1024 pixels. The images in Figure 5A-D were captured in tapping mode at a scan rate of 0.4 Hz and a resolution of 1024 × 1024 pixels. The images in Figure 5EF were captured in tapping mode at a scan rate of 0.5 Hz and a resolution of 4096 x 4096 pixels. The raw images were flattened using Bruker NanoScope Analysis 2.0 software.

### Confocal Raman Spectroscopy

Total protein samples in 10% ethanol were concentrated using Amicon® Ultra Centrifugal 30kDa MWCO regenerated cellulose filters (4mL sample volume, Millipore) to remove any contaminants that would interfere with confocal Raman measurements. 50 µL of the recovered sample was deposited onto a clean, clear mica slide and allowed to dry overnight at room temperature. Confocal Raman measurements were carried out on a LabRam Soleil instrument (Horiba Scientific, France) with 532 nm laser excitation as previously described(71). Raman spectra of the protein rodlets were acquired using 1800 lines/mm grating, 100 µm pinhole, integration time of 60 s with 3 accumulations, and a laser power of 50 mW at the sample plane.

### Thioflavin T assay

ThT assays were made using total protein extracts derived from primary and secondary foam samples, using 40 µM ThT in 50 mM phosphate buffer (pH 7). Prior to sample measurements, total protein extracts, in the dry state, following TFA treatment, were rehydrated with phosphate buffer. Samples were allowed to stand for 30 min at room temperature after a short vortexing step, to allow for rodlet assembly. Right before analysis, an equal amount of ThT was added to the sample and the samples analyzed using a Clario Star microplate UV spectrophotometer. Measurements were carried out using a Greiner 96 F-bottom plate, at room temperature using a 440 nm excitation filter, a 465.5 nm dichroic filter, and a 490 nm filter to detect emission. The setting, scan diameter 3 mm and 16 flashes per well was selected. A focal height of 4.8 mm and 1500 Gain was also selected. Before each cycle, a double orbital mixing mode at 300 rpm (10s) was used. A blank control solution containing only buffer was used for fold change determination (change in fluorescence intensity of the protein sample divided by the fluorescence intensity of the control sample)(60).

### Emulsions and porous Freeze-Dried Emulsion Composites (FDECs)

Emulsions were made by combining 0.8 mg of total proteins extracted from secondary foams with 10% coconut oil (MCT) and 0.3 wt% TEMPO-CNF (zeta potential -50 mV, 5 replicate measurements) in 50 mM phosphate buffer (pH7). The control without protein consisted of the same baseline components. The SDS control consisted of 1% or 0.5% SDS, 10% coconut oil and 0.3 wt% TEMPO-CNF. The emulsions were mixed using a rotor stat homogenizer (Benchmark Scientific D100). Samples were mixed twice for 30s with oil and twice for 20s after TEMPO-CNF addition and then placed at 30°C. For the FDECs, emulsions were prepared as above except 1.9 mg/L total proteins extracted from primary foam was used instead. Samples were immediately frozen at -80°C after mixing for 1 day before freeze-drying for 1 day.

### Scanning Electron Microscopy (SEM) imaging of FDECs

After freeze-drying the FDECs as described above, they were submerged in liquid nitrogen and a small part of each was broken and placed on top of an SEM holder using carbon paste. The sample was coated with a 7 nm layer of platinum (Bal-Tec MED 020 Modular High Vacuum Coating Systems, Bal-Tec AG, Liechtenstein), and assessed via SEM (Qunata 650 FEG ESEM, FEI, Hillsboro, Oregon, USA).

## Acknowledgements

This project was funded by the ETH Board Joint Initiative 2023-2026 Proteins for a Sustainable Future, in the strategic area Energy, Climate and Sustainable Environment. We would like to thank Dr. Rossana Pitocchi for discussions and feedback, Dr. Gilberto Siqueira and Beatriz Arsuffi for preparing the TEMPO oxidized CNF, Dr. Luana Amoroso, Sefora Riillo, and Dr. Svitlana Mykolenko for their help acquiring the food waste for use in experiments and Anja Huch for SEM analysis. The authors gratefully acknowledge the Functional Genomics Center Zurich (FGCZ) of University of Zurich and ETH Zurich, and particularly Dr. Jonas Grossmann, for the Proteomics analyses and technical support.

## References

1. Wösten HAB, Scholtmeijer K. 2015. Applications of hydrophobins: current state and perspectives. Applied Microbiology and Biotechnology 99:1587–1597.

2. Scholtmeijer K, Rink R, Hektor HJ, Wösten HA. 2005. Expression and engineering of fungal hydrophobins. Applied Mycology and Biotechnology 5:239–255.

3. Stringer MA, Timberlake WE. 1995. dewA encodes a fungal hydrophobin component of the Aspergillus spore wall. Molecular microbiology 16:33–44.

4. Talbot NJ, Kershaw MJ, Wakley GE, De Vries O, Wessels JGH, Hamer JE. 1996. MPG1 Encodes a fungal hydrophobin involved in surface interactions during infection-related development of Magnaporthe grisea. Plant Cell 8:985–999.

5. Wessels JGH. 1994. Developmental regulation of fungal cell wall formation. Annual Review of Phytopathology 32:413–437.

6. Seidl-Seiboth V, Gruber S, Sezerman U, Schwecke T, Albayrak A, Neuhof T, Döhren V, Baker S, Kubicek C. 2011. Novel hydrophobins from Trichoderma define a new hydrophobin subclass: protein properties, evolution, regulation and processing. Journal of Molecular Evolution 72:339–351.

7. De Vries O, Fekkes M, Wösten H, Wessels J. 1993. Insoluble hydrophobin complexes in the walls of Schizophyllum commune and other filamentous fungi. Archives of Microbiology 159:330–335.

8. Wessels J, de Vries O, Asgeirsdóttir S, Schuren F. 1991. Hydrophobin genes involved in formation of aerial hyphae and fruit bodies in Schizophyllum. Plant Cell 3:793–799.

9. Scholtmeijer K, De Vocht ML, Rink R, Robillard GT, Wösten HAB. 2009. Assembly of the Fungal SC3 Hydrophobin into Functional Amyloid Fibrils Depends on Its Concentration and Is Promoted by Cell Wall Polysaccharides. Journal of Biological Chemistry 284:26309–26314.

10. Wösten, HAB, de Vocht M. 2000. Hydrophobins, the fungal coat unravelled. Biochimica et Biophysica Acta 1469:79–86.

11. Macindoe I, Kwan AH, Ren Q. 2003. Self-assembly of functional, amphipathic amyloid monolayers by the fungal hydrophobin EAS. Proceedings of the National Academy of Sciences of the United States of America 109:E804–E811.

12. Stroud PA, Goodwin JS, Butko P, Cannon GC, McCormick CL. 2003. Experimental evidence for multiple assembled states of Sc3 from Schizophyllum commune. Biomacromolecules 4:956–967.

13. Gravagnuolo AM, Longobardi S, Luchini A, Appavou MS, De Stefano L, Notomista E, Paduano L, Giardina P. 2016. Class I hydrophobin Vmh2 adopts atypical mechanisms to self-assemble into functional amyloid fibrils. Biomacromolecules 17:954–964.

14. Xu D, Wang Y, Keerio AA, Ma A. 2021. Identification of hydrophobin genes and their physiological functions related to growth and development in Pleurotus ostreatus. Microbiological Research 247.

15. Karlsson M, Stenlid J, Olson Å. 2007. Two hydrophobin genes from the conifer pathogen Heterobasidion annosum are expressed in aerial hyphae. Mycologia 99:227–231.

16. Tasaki Y, Ohata K, Hara T, Joh T. 2004. Three genes specifically expressed during phosphate deficiency in Pholiota nameko strain N2 encode hydrophobins. Current Genetics 45:19–27.

17. Pothiratana C, Fuangsawat W, Jintapattanakit A, Teerapatsakul C, Thachepan S. 2021. Putative hydrophobins of black poplar mushroom (Agrocybe cylindracea). Mycology 12:58–67.

18. MJ K, NJ T. 1998. Hydrophobins and repellents: proteins with fundamental roles in fungal morphogenesis. Fungal Genetics and Biology 23:18–33.

19. Taylor JW, Spatafora J, O Donnell K, Lutzoni F, James T, Hibbett DS, Geiser D, Bruns TD, Blackwell M. 2004. The Fungi, p. 171–194. *In* Cracraft, J. Donoghur, M.J. (ed.), Assembling the Tree of Life. Oxford University Press, New York.

20. Qiao J, Liu H, Xue P, Hong M, Guo X, Xing Z, Zhao M, Zhu J. 2023. Function of a hydrophobin in growth and development, nitrogen regulation, and abiotic stress resistance of Ganoderma lucidum. FEMS Microbiology Letters 370:1–10.

21. Devi MsKS, S Sundareshwar. 2023. Food Waste Management System. IJRASET 11:2462–2466.

22. Porter SD, Reay DS, Higgins P, Bomberg E. 2016. A half-century of production-phase greenhouse gas emissions from food loss & waste in the global food supply chain. Science of The Total Environment 571:721–729.

23. Ramandani AA, Sun Y, Lan JC, Lim JW, Chang J, Srinuanpan S, Khoo KS. 2024. Upcycling food waste as a low-cost cultivation medium for *Chlorella* sp. microalgae. J Sci Food Agric jsfa.13910.

24. Escaramboni B, Garnica BC, Abe MM, Palmieri DA, Fernández Núñez EG, De Oliva Neto P. 2022. Food Waste as a Feedstock for Fungal Biosynthesis of Amylases and Proteases. Waste Biomass Valor 13:213–226.

25. Kulkarni SS, Nene SN, Joshi KS. 2020. Exploring malted barley waste for fungi producing surface active proteins like hydrophobins. SN Appl Sci 2.

26. Kulkarni SS, Nene SN, Joshi KS. 2020. A comparative study of production of hydrophobin like proteins (HYD-LPs) in submerged liquid and solid state fermentation from white rot fungus Pleurotus ostreatus. Biocatalysis and Agricultural Biotechnology 23:101440.

27. Kelly JM, Katz ME. 2010. Glucose, p. 291–311. *In* Cellular and Molecular Biology of Filamentous Fungi. ASM Press, Washington DC.

28. Chroumpi T, Mäkelä M, de Vries R. 2020. Engineering of primary carbon metabolism in filamentous fungi. Biotechnology Advances 43.

29. Liu L, Feng J, Gao K, Zhou S, Yan M, Chuanhong T, Zhou J, Liu Y, Zhang J. 2022. Influence of carbon and nitrogen sources on structural features and immunomodulatory activity of exopolysaccharides from Ganoderma lucidum. Process Biochemistry 119:96–105.

30. Fraga I, Coutinho J, Bezerra RM, Dias AA, Marques G, Nunes FM. 2014. Influence of culture medium growth variables on Ganoderma lucidum exopolysaccharides structural features. Carbohydrate Polymers 111:936–946.

31. Ho P-Y, Namasivayam P, Sundram S, Ho C-L. 2020. Expression of genes encoding manganese peroxidase and laccase of Ganoderma boninense in response to nitrogen sources, hydrogen peroxide and phytohormones. Genes 11:1263.

32. Lee K-M, Lee S-Y, Lee H-Y. 1999. Effect of ammonium phosphate on mycelial growth and exopolysaccharides production of Ganoderma lucidum in an air-lift fermenter. Journal of microbiology and biotechnology 9:726–731.

33. Maski S, Ngom SI, Rached B, Chouati T, Benabdelkhalek M, El Fahime E, Amar M, Béra-Maillet C. 2021. Hemicellulosic biomass conversion by Moroccan hot spring Bacillus paralicheniformis CCMM B940 evidenced by glycoside hydrolase activities and whole genome sequencing. Biotech 11:1–13.

34. Fotirić Akšić M, Dabić Zagorac D, Gašić U, Tosti T, Natić M, Meland M. 2022. Analysis of apple fruit (Malusx domestica Borkh) quality attriubtes obtained from organic and integrated production systems. Sustainability 14:5300.

35. Salem MM, Ibrahim S, Kim C, Seo CW, Shahbazi A, AbuGayaleh A. 2009. Lactic acid production from apple skin waste by immobilized cells of Lactobacillus reuteri, p. 31–37. *In* Proceedings of the 2007 National Conference on Environmental Science and Technology. Springer New York.

36. Monago-Maraña O, Afseth NK, Knutsen SH, Wubshet SG, Wold JP. 2021. Quantification of soluble solids and individual sugars in apples by Raman spectroscopy: A feasibility study. Postharvest Biology and Technology 180.

37. Gupta E, Mishra P, Shiekh A, Gupta K. 2020. Fruit peels: a strong natural source of antioxidant and prebiotics. Carpathian Journal of Food Science & Technology 12.

38. Yajing L, Sun H, Li J, Qin S, Yang W, Ma X, Qiao X, Yang B. 2021. Effects of genetic background and altitude on sugars, malic acid and ascorbic acid in fruits of wild and cultivated apples (Malus sp.). Foods 10:2950.

39. Arnous A, Meyer AS. 2009. Quantitative prediction of cell wall polysaccharide composition in grape (Vitis vinifera L.) and apple (Malus domestica) skins from acid hydrolysis monosaccharide profiles. Journal of agricultural and food chemistry 57:3611–3619.

40. Tomé D. 2021. Yeast extracts: nutritional and flavoring food ingredients. ACS Food Science and Technology 1:487–494.

41. Selvaraj B, Sanjeevirayar A, Rajendran A. 2015. Laccase production using mixed substrates containing lignocellulosic materials by Pleurotus ostreatus in submerged liquid culture. International Journal of ChemTech Research 7:355–368.

42. Aydınoğlu T, Sargın S. 2013. Production of laccase from Trametes versicolor by solidstate fermentation using olive leaves as a phenolic substrate. Bioprocess and biosystems engineering, 36:215–222.

43. Giovani G, Rosi I. 2007. Release of cell wall polysaccharides from Saccharomyces cerevisiae thermosensitive autolytic mutants during alcoholic fermentation. International Journal of Food Microbiology 116:19–24.

44. Blasco L, Viñas M, Villa TG. 2011. Proteins influencing foam formation in wine and beer: The role of yeast. International Microbiology 14:61–71.

45. Schönig B, Brown DW, Oeser B, Tudzynski B. 2008. Cross-species hybridization with Fusarium verticillioides microarrays reveals new insights into Fusarium fujikuroi nitrogen regulation and the role of AreA and NMR. Eukaryotic Cell 7:1831–1846.

46. Macios M, Caddick MX, Weglenski P, Scazzocchio C, Dzikowska A. 2012. The GATA factors AREA and AREB together with the co-repressor NMRA, negatively regulate arginine catabolism in Aspergillus nidulans in response to nitrogen and carbon source. Fungal Genetics and Biology 49:189–198.

47. Neuhof T, Dieckmann R, Druzhinina IS, Kubicek CP, Nakari-Setälä T, Penttilä M, Von Döhren H. 2007. Direct identification of hydrophobins and their processing in Trichoderma using intact-cell MALDI-TOF MS. FEBS Journal 274:841–852.

48. Nakari-Setälä T, Aro N, Kalkkinen N, Alatalo E, Penttilä M. 1996. Genetic and biochemical characterization of the Trichoderma reesei hydrophobin HFBI. European Journal of Biochemistry 235:248–255.

49. Nakari-Setälä T, Aro N, Ilmén M, Muñoz G, Kalkkinen N, Penttilä M. 1996. Differential expression of the vegetative and spore-bound hydrophobins of Trichoderma reesei: Cloning and characterization of the hfb2 gene. VTT Publications 423:1–15.

50. Vergunst KL, Langelaan DN. 2022. The N-terminal tail of the hydrophobin SC16 is not required for rodlet formation. Scientific Reports 12.

51. Morris V, Kwan A, Sunde M. 2013. Analysis of the structure and conformational states of DewA gives insight into the assembly of the fungal hydrophobins. Journal of Molecular Biology 425:244–56.

52. Rea I, Giardina P, Longobardi S, Porro F, Casuscelli V, Rendina I, De Stefano L. 2012. Hydrophobin Vmh2–glucose complexes self-assemble in nanometric biofilms. Journal of the Royal Society Interface 9:2450–2456.

53. Usov I, Nyström G, Adamcik J, Handschin S, Schütz C, Fall A, Bergström L, Mezzenga R. 2015. Understanding nanocellulose chirality and structure-properties relationship at the single fibril level. Nature Communications 6.

54. Žganec M, Taler Verčič A, Muševič I, Škarabot M, Žerovnik E. 2023. Amyloid fibrils of stefin B show anisotropic properties. International Journal of Molecular Sciences 24.

55. Usov I, Mezzenga R. 2015. FiberApp: An Open-Source Software for Tracking and Analyzing Polymers, Filaments, Biomacromolecules, and Fibrous Objects.

56. Sane SU, Cramer SM, Przybycien TM. 1999. A Holistic Approach to Protein Secondary Structure Characterization Using Amide I Band Raman Spectroscopy. Analytical Biochemistry 269:255–272.

57. Sarroukh R, Goormaghtigh E, Ruysschaert J-M, Raussens V. 2013. ATR-FTIR: A “rejuvenated” tool to investigate amyloid proteins. Biochimica et Biophysica Acta (BBA) Biomembranes 1828:2328–2338.

58. Maiti NC, Apetri MM, Zagorski MG, Carey PR, Anderson VE. 2004. Raman Spectroscopic Characterization of Secondary Structure in Natively Unfolded Proteins: α-Synuclein. J Am Chem Soc 126:2399–2408.

59. De Vocht ML, Reviakine I, Ulrich W, Bergsma-Schutter W, Wösten HAB, Vogel H, Brisson A, Wessels JGH, Robillard GT. 2002. Self-assembly of the hydrophobin SC3 proceeds via two structural intermediates. Protein Science 11:1199–1205.

60. Xue C, Lin TY, Chang D, Guo Z. 2017. Thioflavin T as an amyloid dye: fibril quantification, optimal concentration and effect on aggregation. R Soc open sci 4:160696.

61. Jeon J, Yau W-M, Tycko R. 2023. Early events in amyloid-β self-assembly probed by time-resolved solid state NMR and light scattering. Nat Commun 14:2964.

62. Girych M, Gorbenko G, Maliyov I, Trusova V, Mizuguchi C, Saito H, Kinnunen P. 2016. Combined thioflavin T–Congo red fluorescence assay for amyloid fibril detection. Methods Appl Fluoresc 4:034010.

63. Robbins KJ, Liu G, Lin G, Lazo ND. 2011. Detection of Strongly Bound Thioflavin T Species in Amyloid Fibrils by Ligand-Detected^1^ H NMR. J Phys Chem Lett 2:735–740.

64. Han J, Kawauchi M, Terauchi Y, Yoshimi A, Tanaka C, Nakazawa T, Honda Y. 2023. Physiological function of hydrophobin Vmh3 in lignin degradation by white-rot fungus *Pleurotus ostreatus*. Letters in Applied Microbiology 76:ovad048.

65. Wu H, Nakazawa T, Xu H, Yang R, Bao D, Kawauchi M, Sakamoto M, Honda Y. 2021. Comparative transcriptional analyses of Pleurotus ostreatus mutants on beech wood and rice straw shed light on substrate-biased gene regulation. Appl Microbiol Biotechnol 105:1175–1190.

66. Eslami E, Donsì F, Ferrari G, Pataro G. 2024. Enhancing Cutin Extraction Efficiency from Industrially Derived Tomato Processing Residues by High-Pressure Homogenization. Foods 13:1415.

67. Leonowicz A, Matuszewska A, Luterek J, Ziegenhagen D, Wojtaś-Wasilewska M, Cho NS, Hofrichter M, Rogalski J. 1999. Biodegradation of lignin by white rot fungi. Fungal Genetics and Biology 27:175–185.

68. Dons JJM, De Vries OMH, Wessels JGH. 1979. Characterization of the genome of the basidiomycete Schizophyllum commune. BBA Section Nucleic Acids And Protein Synthesis 563:100–112.

69. Morris VK, Ren Q, Macindoe I, Kwan AH, Byrne N, Sunde M. 2011. Recruitment of class I hydrophobins to the air:Water interface initiates a multi-step process of functional amyloid formation. Journal of Biological Chemistry 286:15955–15963.

70. Reyes C, Ahrendt S, Riley R, Lipzen A, Ng V, Grigoriev IV, Schwarze FWMR, Baars O. 2025. Siderophores and secondary metabolites produced by Ganoderma adspersum. Microbiology 171.

71. Mrđenović D, Combes BF, Ni R, Zenobi R, Kumar N. 2024. Probing Chemical Complexity of Amyloid Plaques in Alzheimer’s Disease Mice using Hyperspectral Raman Imaging. ACS Chem Neurosci 15:78–85.

