## Supporting Information for "Fungal hydrophobins unleashed from food waste: production, rodlet assembly, and functional properties in *Ganoderma adspersum*"

**Table S1. Annotated sugar uptake transporters in *Ganoderma adspersum*.** Transporter genes predicted to facilitate uptake of glucose, galactose, and xylose were identified in the genome of *G. adspersum* using the MycoCosm annotation pipeline. Gene IDs, and annotation sources are listed.

| **Protein ID(s)** | **MycoCosm Annotations** |
| --- | --- |
| 566641<br>581557 | 2.A.1.1.68The Glucose Transporter/Sensor Rgt2 |
| 891635 | 2.A.1.1.36 The low affinity, glucose-inducible glucose transporter, MstE (Forment et al., 2006[1]) |
| 260246 | 2.A.7.9.3 Chloroplast phosphoenolpyruvate:Pi antiporter (PPT) (triose-Ps and glycerate- Ps are poor substrates), KOG1441Glucose-6-phosphate/phosphate and phosphoenolpyruvate/phosphate antiporter |
| 538919, 557954<br>637653, 649227<br>757878,779654<br>872320,874934 | 2.A.1.7.2Glucose/galactose porter |
| 663919<br>734311 | 2.A.1.7.3 Glucose/Mannose/Xylose: H+ symporter (Paulsen et al., 1998[2]; G.Gosset, personal communication). |
| 724361 | KOG1441Glucose-6-phosphate/phosphate and phosphoenolpyruvate/phosphate antiporter, 2.A.7.9.1Chloroplast triose-P/glycerate-3-P:Pi antiporter (TPT) (phosphoenolpyruvate and 2-phosphoglycerate are poor substrates). |
| 728060 | KOG1441Glucose-6-phosphate/phosphate and phosphoenolpyruvate/phosphate antiporter, 2.A.7.9.4Sly41p (transport function unknown) |
| 547798 | KOG0255 Synaptic vesicle transporter SVOP and related transporters (major facilitator superfamily), 2.A.1.2.46 The fructose/glucose uniporter, Ffz2 (64% identical to 2.A.1.2.23). Both sugars are transported with similar affinities and efficiencies (Leandro et al., 2011[3]). |
| 555237 | 2.A.1.1.6 Galactose, glucose uniporter, Gal2 |
| 632861 | 2.A.1.1.39 The high affinity glucose transporter, Hgt1 (Baruffini et al., 2006[4]) |
| 637757 | 2.A.1.1.40 The xylose facilitator, Xylhp (Nobre et al., 1999[5]) |
| 696061 | 2.A.1.1.43 The monosaccharide (MST) (glucose > mannose > galactose > fructose):H+ symporter, MST1 (Schussler et al., 2006[6]). |
| 738271<br>818310 | 2.A.1.1.69 Sugar & polyol transporter 1 (SPT1): broad specificity; takes up glucose (Schilling and Oesterhelt, 2007[7]) |
| 758642 | 2.A.1.1.38 The glycerol:H+ symporter, Stl1p (Ferreira et al., 2005) |
| 879916 | 2.A.1.1.57 High affinity (15 µM) glucose (monosaccharides including xylose):H+ symporter, MstA (Jørgensen et al., 2007[8]). |
| 681820 | 2.A.7.22.2 The undecaprenyl phosphate-?-aminoarabinose flippase ArnE/ArnF heterodimer from the cytoplasm to the periplasm (Yan et al., 2007[9]). |
| 901165 | KOG0369 Pyruvate carboxylase, 3.B.1.1.1Na+-transporting oxalo-acetate decarboxylase |

**Table S2. Annotated arabinose-related genes in *Ganoderma adspersum*.** Genes associated with arabinose metabolism and export were identified in the genome of *G. adspersum*. No arabinose uptake transporters were annotated.

| **Protein ID(s)** | **MycoCosm Annotations** |
|---|---|
| 750175 | 1.1.1.116D-arabinose 1-dehydrogenase (NAD+) |
| 873307,<br>757292 | The acriflavin-sensitivity protein, YnfM (increases sensitivity to acriflavin specifi-cally). Also exports arabinose but not xylose (Koita and Rao 2012[10]). |
| 773129 | 3.2.1.55non-reducing end alpha-L-arabino furanosidase |
| 573047 | KOG2517Ribulose kinase and related carbohydrate kinases |
| 483776<br>884918 | KOG3111D-ribulose-5-phosphate 3-epimerase |

**Table S3.** Carbon and nitrogen content of food wastes used in this study

| Food waste | Carbon | Nitrogen | Comments | References |
|---|---|---|---|---|
| **Apple skins** | 47.71 ± 0.37 % | 1.10 ± 0.02 % | Measurements based on CHN micro elemental analyzer | This study |
| **Flaxseed (brown)** | 45.33 ± 0.05 % | 5.31 ± 0.07 % | Measurements based on CHN micro elemental analyzer | This study |
| **Flaxseed (golden)** | 45.33 ± 0.17 % | 6.95 ± 0.23 % | Measurements based on CHN micro elemental analyzer | This study |
| **Barley flakes** | 48.30 ± 0.43 % | 3.60 ± 0.08 % | Measurements based on CHN micro elemental analyzer | This study |
| **Media** | **Carbon** | **Nitrogen** | **Comments** | **References** |
| **SCMM (per L)** | 32.36 %C of dry solutes (incl. buffer) | 1.07 %N of dry solutes (incl. buffer) | Based on % of dry solutes in the medium | Dons et al.1979 [11] |

* Calculated for 22 g $L^{-1}$ glucose·$H_2O$ and 1.5 g $L^{-1}$ L-asparagine·$H_2O$; includes 0.5 g $L^{-1}$ $MgSO_4 \cdot 7H_2O$, trace elements, $FeCl_3 \cdot 6H_2O$, thiamine, and the phosphate buffer salts (0.551 g $KH_2PO_4$ + 1.568 g $K_2HPO_4$ per L) in the % denominator. Salts/stocks contribute no carbon (and only negligible nitrogen); C and N loads are unchanged by including the buffer in the denominator.

**Table S4.** Comparison of secondary structural information of Class I hydrophobins based on FTIR results of SC3[12,13] and Raman Confocal Microscopy determined in this study.

| FTIR Analysis | | | Raman Spectroscopy | Raman Spectroscopy |
|---|---|---|---|---|
| | *S. commune* SC3 (soluble) (O/N drying on surface)* | *S. commune* SC3 (air/water) (O/N drying on surface)** | 1° foam aggregates (non-food waste) | 2° foam aggregates (food waste) |
| **β-sheet** | 41% | 65% | 66% | 65% |
| **Random Coil** | 20% | 10% | 4% | 5.6% |
| **α-helix** | 23% | 16% | 29% | 29 |
| **β-turn** | 16% | 9% | nm | nm |

***SC3 (soluble):** de Vocht, M. L., Scholtmeijer, K., et al. (1998).: https://doi.org/10.1016/S0006-3495(98)77912-3

**Zangi, Ronen, et al. (2002) 10.1016/S0006-3495(02)75153-9

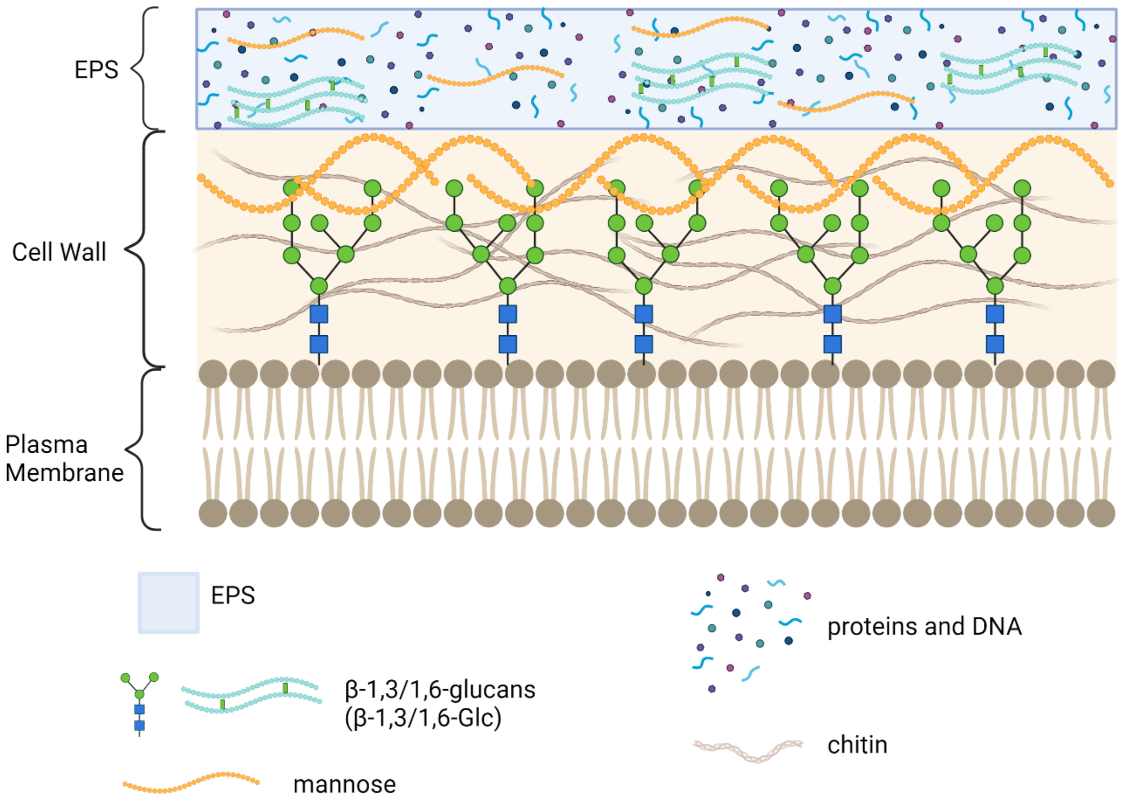


**Figure S1**. Schematic of fungal cell wall with sugars, chitin and EPS. Image generated with BioRender.

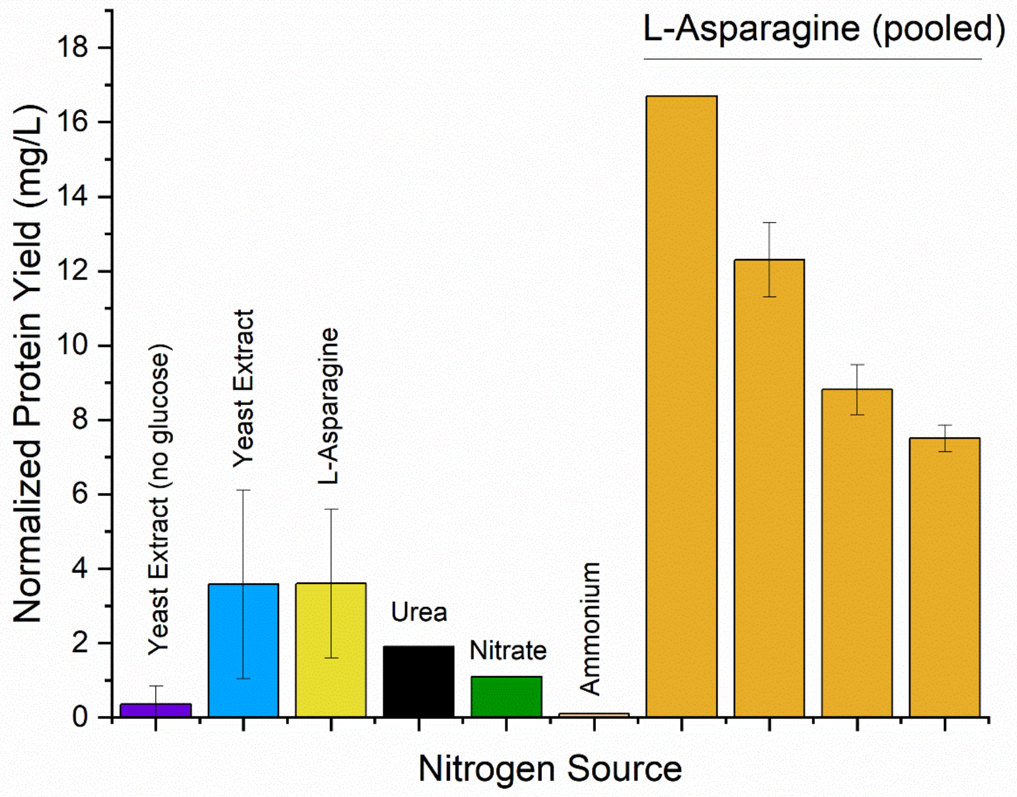


**Figure S2**. Normalized protein yield in primary foam from 1L of SCMM minimal media supplemented with various nitrogen sources. Pooled samples from different experiments are indicated as four different columns.

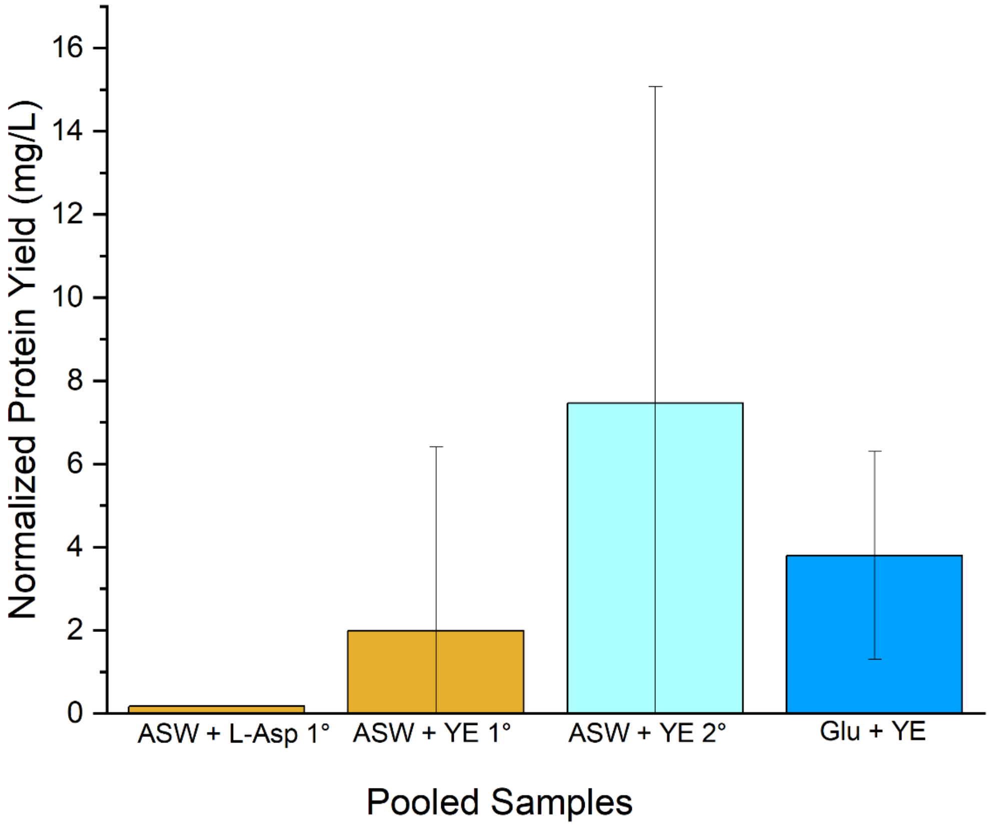


**Figure S3.** Normalized protein yield in primary (1°) and secondary (2°) foam from 1L minimal media supplemented with apple skin waste. No glucose was present in the media unless indicated. If L-asparagine was present (instead of yeast extract (YE) it is indicated as L-Asp. Pooled samples from experiments are indicated as pooled.

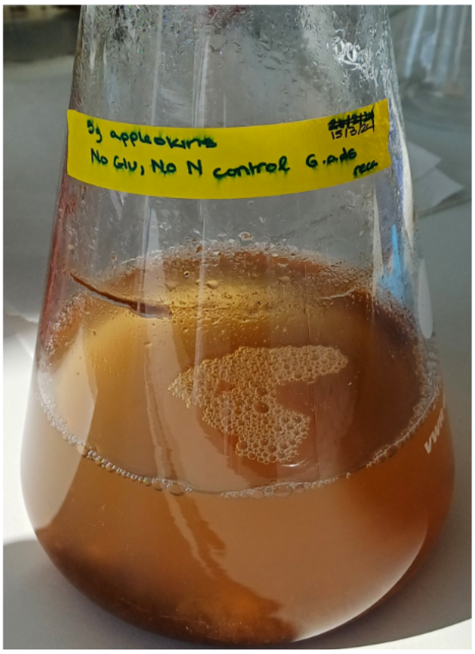


**Figure S4.** *G. adspersum* culture containing 5 g of apple skin waste with no added glucose or nitrogen source.

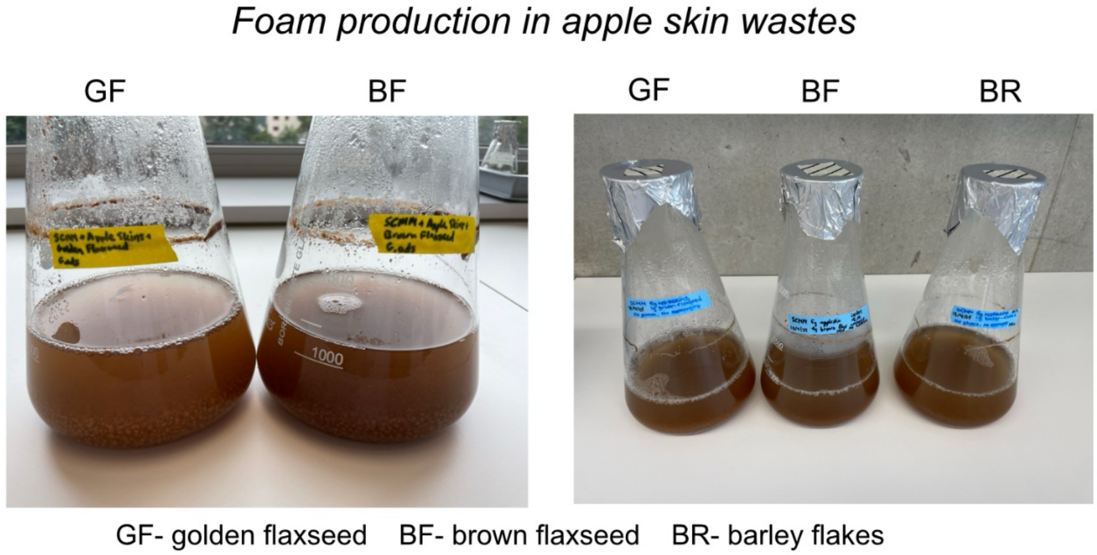


**Figure S5**. Amount of primary foam formed after 12 days of growth in *G. adspersum* cultures containing 5g of apple skin waste and flaxseed or barley flakes.

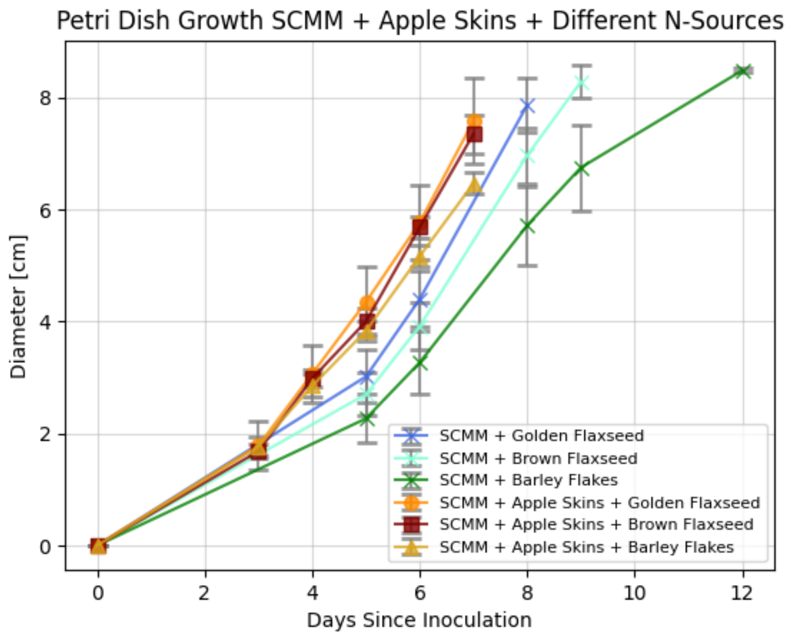


**Figure S6**. Growth rate of *G. adspersum* in the presence of SCMM and various food wastes on solid Petri dishes. No additional nitrogen source was added to SCMM + golden flaxseed, SCMM + brown flaxseed and SCMM + barley flakes (glucose was present). No glucose or additional nitrogen source was added to SCMM + apple skins containing flaxseed or barley flakes.

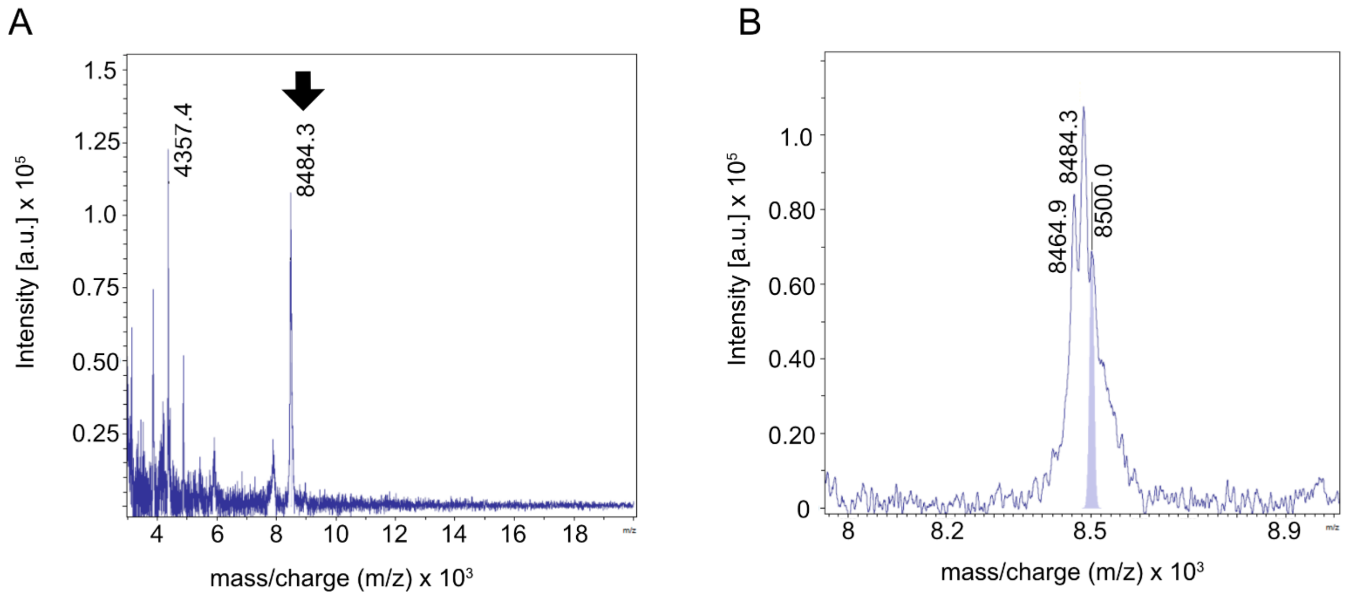


**Figure S7**. Mass spectra of signals detected in secondary foam samples from *G.adspersum* grown with apple skins and yeast extract. **A)** Mass range of hydrophobin proteins of interest. **B)** Close-up of the mass spectra in A showing three individual peaks corresponding to hydrophobins.

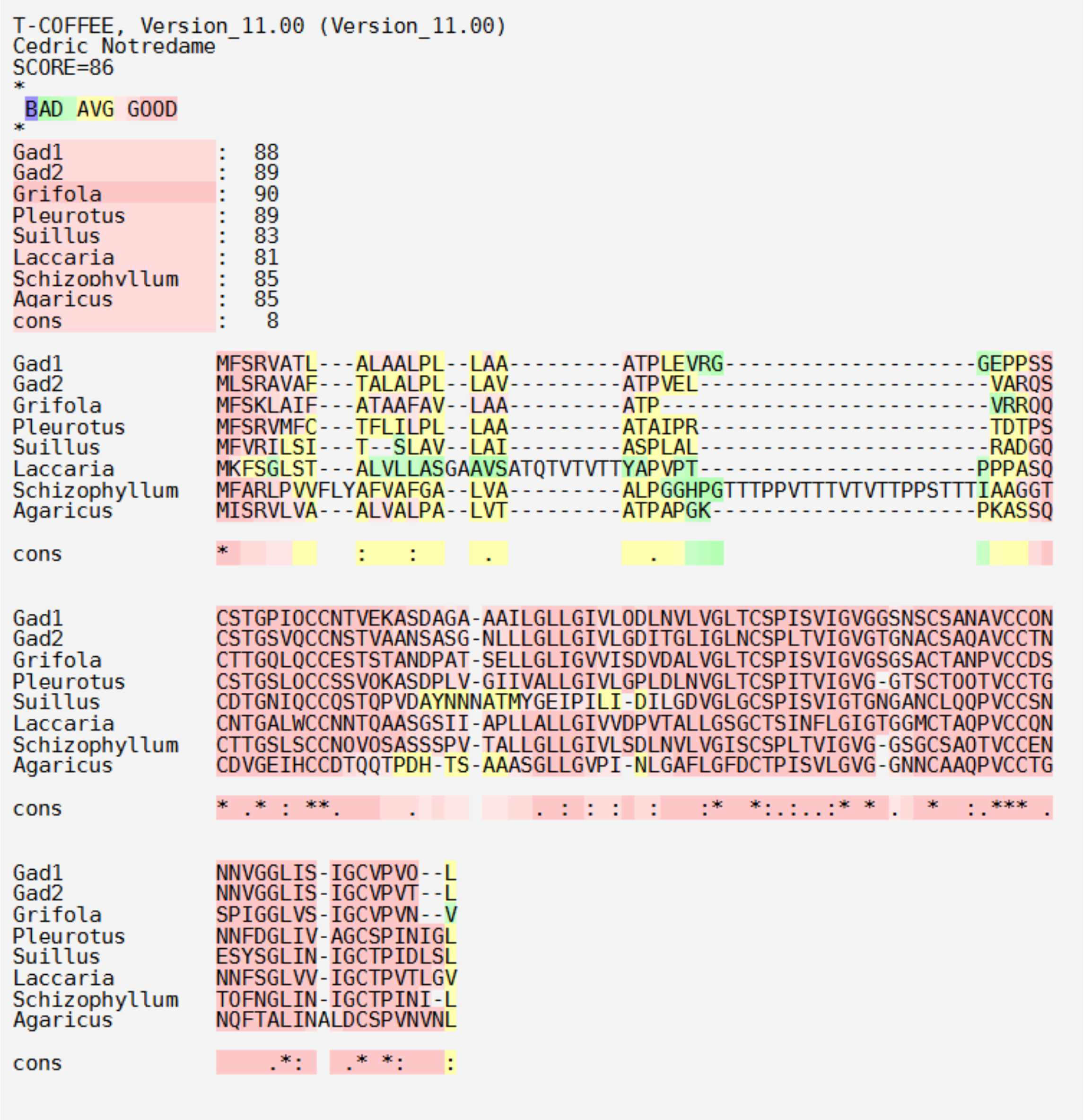


**Figure S8.** Amino acid alignment of class I hydrophobin sequences Gad1 and Gad2 from *G. adspersum* with other fungal class I hydrophobin sequences[14] including: HGFI, *Grifola frondosa* (accession no. ABO42329), VMH2; *Pleurotus ostreatus* PC15 (accession no. KDQ22759), LbH1; *Laccaria bicolor* S238N-H82 (accession no. DS547093.1), Hyd3; *Schizophillum commune* (accession no. P16933) and ABH1; *Agaricus bisporus* (accession no. P49072). Residues in pink indicate high similarity between proteins. Conserved residues, including cysteine (C), are marked with an asterisk (*). Sequences were aligned using the default input setting of T-coffee[15–19]. Sequences were retrieved from the National Center for Biotechnology Information website. https://www.ncbi.nlm.nih.gov/

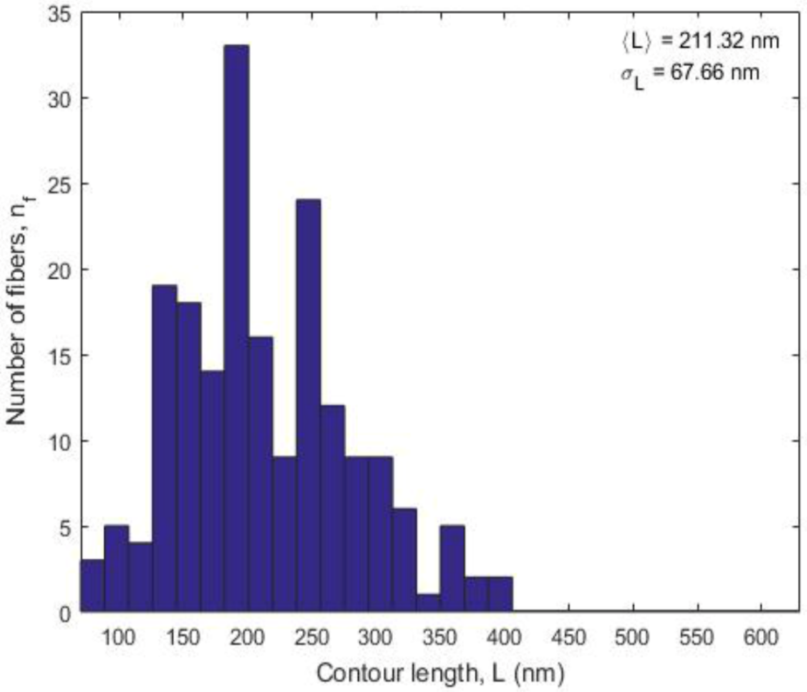


**Figure S9**. Histogram of rodlet contour lengths measured from AFM images (n= 190 fibers, analyzed with FiberApp)[20]. The average length was 211 ± 68 nm, indicating a heterogeneous distribution with most rodlets falling between ~140-280 nm. A smaller fraction of longer fibers up to ~450 nm was also observed.

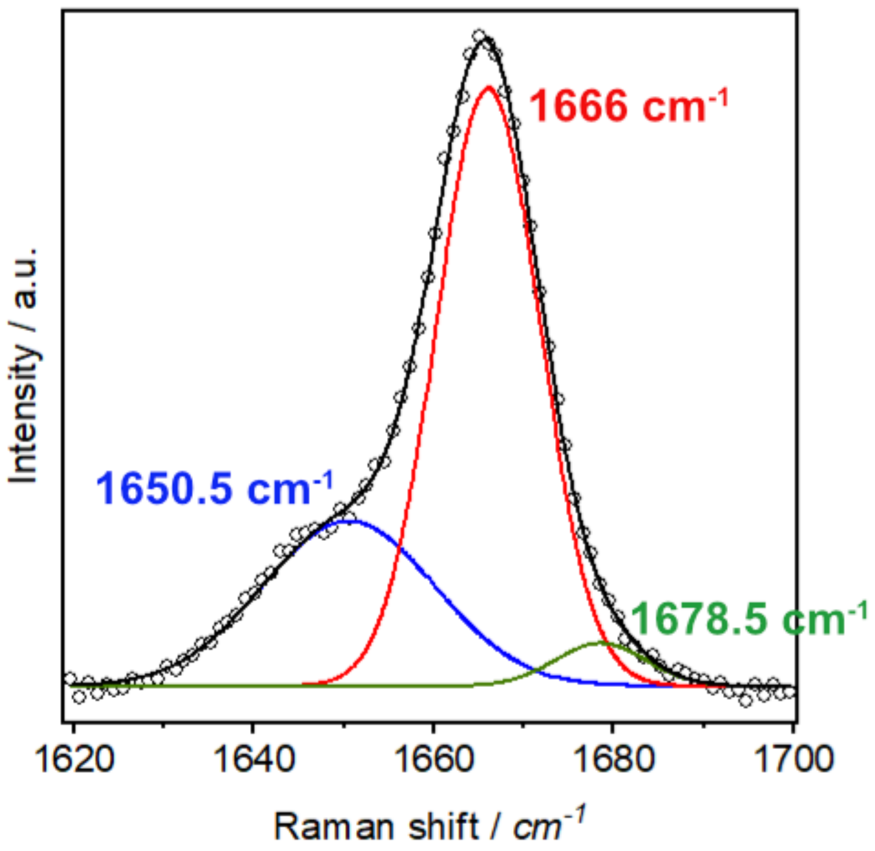


**Figure S10.** Representative deconvolution of the Amide I band, from foams produced in non-food waste medium, showing Gaussian fitting into β-sheet, α-helix, and coil/polyproline II components.

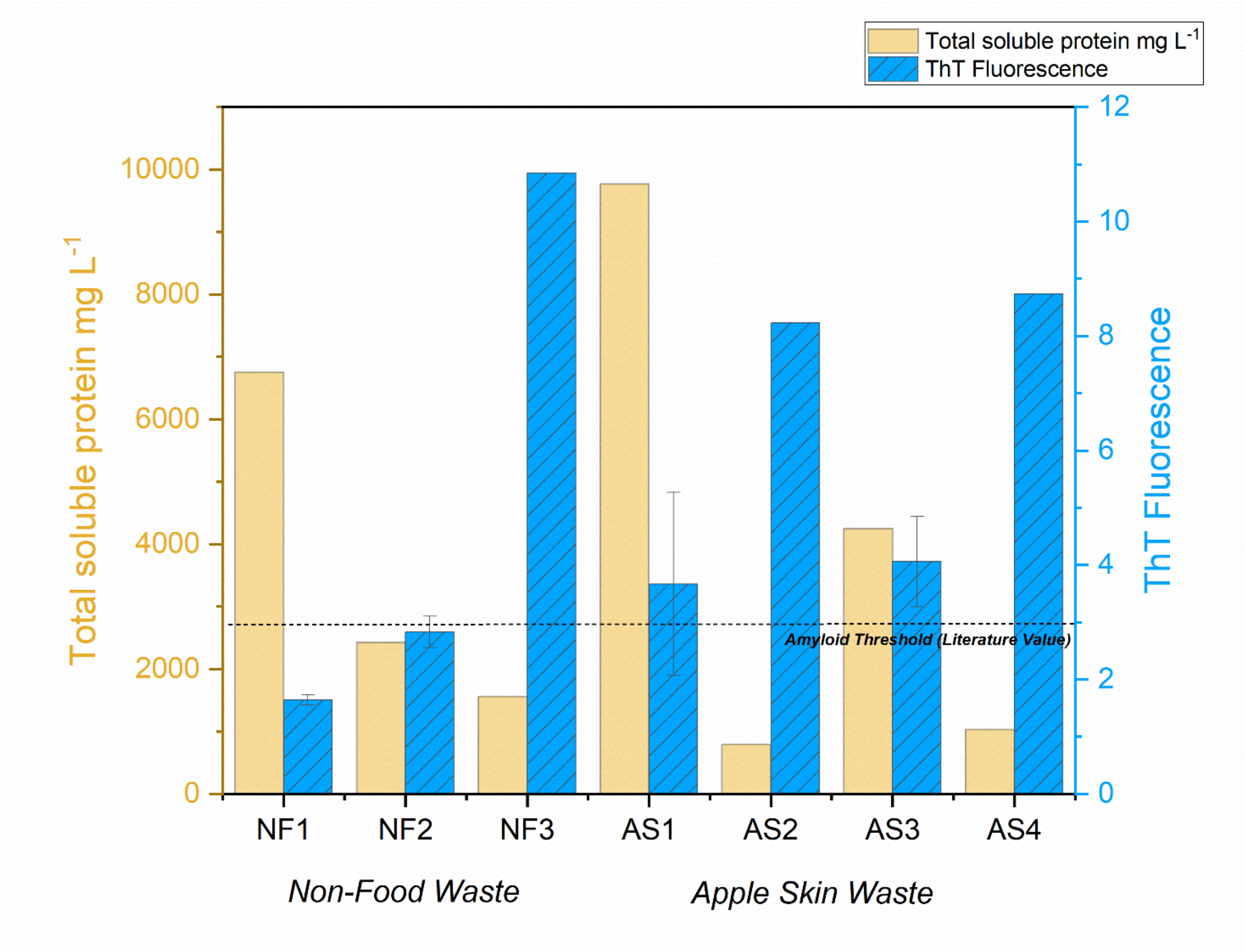


**Figure S11**. **Total soluble protein concentration and ThT fluorescence of foam derived from cultures grown with and without apple skin waste.** Bars show protein concentration (left axis, orange) and ThT fluorescence (right axis, blue) for foam derived from cultures grown without food waste (NF) and for foam derived from cultures grown with apple skin waste (AS). Numbers following NF and AS labels refer to protein extracts from different experiments. Standard deviation bars are from replicate samples. The dotted line indicates the literature defined threshold for significant amyloid-like aggregation[21,22].

### Supporting References


1. Forment, J. V., Flipphi, M., Ramón, D., Ventura, L. & MacCabe, A. P. Identification of the mstE Gene Encoding a Glucose-inducible, Low Affinity Glucose Transporter in Aspergillus nidulans*. *Journal of Biological Chemistry* **281**, 8339–8346 (2006).
2. Paulsen, I. T., Chauvaux, S., Choi, P. & Saier, M. H. Characterization of Glucose-Specific Catabolite Repression-Resistant Mutants of *Bacillus subtilis*: Identification of a Novel Hexose:$H^+$ Symporter. *Journal of Bacteriology* **180**, 498–504 (1998).
3. Leandro, M. J., Sychrová, H., Prista, C. & Loureiro-Dias, M. C. The osmotolerant fructophilic yeast Zygosaccharomyces rouxii employs two plasma-membrane fructose uptake systems belonging to a new family of yeast sugar transporters. *Microbiology* vol. 157 601–608 (2011).
4. Baruffini, E., Goffrini, P., Donnini, C. & Lodi, T. Galactose transport in Kluyveromyces lactis: major role of the glucose permease Hgt1. *FEMS Yeast Research* **6**, 1235–1242 (2006).
5. Nobre, A., Lucas, C. & Leão, C. Transport and Utilization of Hexoses and Pentoses in the Halotolerant Yeast *Debaryomyces hansenii*. *Applied and Environmental Microbiology* **65**, 3594–3598 (1999).
6. Schüßler, A., Martin, H., Cohen, D., Fitz, M. & Wipf, D. Characterization of a carbohydrate transporter from symbiotic glomeromycotan fungi. *Nature* **444**, 933–936 (2006).
7. Schilling, S. & Oesterhelt, C. Structurally reduced monosaccharide transporters in an evolutionarily conserved red alga. *Biochem J* **406**, 325–331 (2007).
8. Jørgensen, T. R. *et al.* Glucose uptake and growth of glucose-limited chemostat cultures of Aspergillus niger and a disruptant lacking MstA, a high-affinity glucose transporter. *Microbiology* vol. 153 1963–1973 (2007).
9. Yan, A., Guan, Z. & Raetz, C. R. H. An Undecaprenyl Phosphate-Aminoarabinose Flippase Required for Polymyxin Resistance in Escherichia coli*. *Journal of Biological Chemistry* **282**, 36077–36089 (2007).
10. Koita, K. & Rao, C. V. Identification and Analysis of the Putative Pentose Sugar Efflux Transporters in Escherichia coli. *PLOS ONE* **7**, 1–10 (2012).
11. Dons, J. J. M., De Vries, O. M. H. & Wessels, J. G. H. Characterization of the genome of the basidiomycete Schizophyllum commune. *BBA Section Nucleic Acids And Protein Synthesis* **563**, 100–112 (1979).
12. De Vocht, M. L. *et al.* Self-assembly of the hydrophobin SC3 proceeds via two structural intermediates. *Protein Science* **11**, 1199–1205 (2002).
13. Zangi, R., De Vocht, M. L., Robillard, G. T. & Mark, A. E. Molecular Dynamics Study of the Folding of Hydrophobin SC3 at a Hydrophilic/Hydrophobic Interface. *Biophysical Journal* **83**, 112–124 (2002).
14. Pothiratana, C., Fuangsawat, W., Jintapattanakit, A., Teerapatsakul, C. & Thachepan, S. Putative hydrophobins of black poplar mushroom (Agrocybe cylindracea). *Mycology* **12**, 58–67 (2021).
15. Notredame, C., Higgins, D. G. & Heringa, J. T-coffee: A novel method for fast and accurate multiple sequence alignment. *Journal of Molecular Biology* **302**, 205–217 (2000).
16. Di Tommaso, P. *et al.* T-Coffee: A web server for the multiple sequence alignment of protein and RNA sequences using structural information and homology extension. *Nucleic Acids Research* **39**, 13–17 (2011).
17. Armougom, F. *et al.* Expresso: Automatic incorporation of structural information in multiple sequence alignments using 3D-Coffee. *Nucleic Acids Research* **34**, 604–608 (2006).
18. O'Sullivan, O., Suhre, K., Abergel, C., Higgins, D. G. & Notredame, C. 3DCoffee: Combining protein sequences and structures within multiple sequence alignments. *Journal of Molecular Biology* **340**, 385–395 (2004).
19. Poirot, O., Suhre, K., Abergel, C., O'Toole, E. & Notredame, C. 3DCoffeeιgs: A web server for combining sequences and structures into a multiple sequence alignment. *Nucleic Acids Research* **32**, 37–40 (2004).
20. Usov, I. & Mezzenga, R. FiberApp: An Open-Source Software for Tracking and Analyzing Polymers, Filaments, Biomacromolecules, and Fibrous Objects. (2015).
21. Girych, M. *et al.* Combined thioflavin T–Congo red fluorescence assay for amyloid fibril detection. *Methods Appl. Fluoresc.* **4**, 034010 (2016).
22. Robbins, K. J., Liu, G., Lin, G. & Lazo, N. D. Detection of Strongly Bound Thioflavin T Species in Amyloid Fibrils by Ligand-Detected[1] H NMR. *J. Phys. Chem. Lett.* **2**, 735–740 (2011).
